# Public LC-Orbitrap-MS/MS Spectral Library for Metabolite Identification

**DOI:** 10.1101/2020.11.21.392266

**Authors:** Prasad Phapale, Andrew Palmer, Rose Muthoni Gathungu, Dipali Kale, Britta Brügger, Theodore Alexandrov

## Abstract

Liquid chromatography-mass spectrometry(LC-MS)-based untargeted metabolomics studies require high-quality spectral libraries for reliable metabolite identification. We have constructed EMBL-MCF, an open LC-MS/MS spectral library that currently contains over 1600 fragmentation spectra from 435 authentic standards of endogenous metabolites and lipids. The unique features of the library are presence of chromatographic profiles acquired with different LC-MS methods and coverage of different adduct ions. The library covers many biologically important metabolites with some unique metabolites and lipids as compared to other public libraries. The EMBL-MCF spectral library is created and shared using an *in-house* developed web-application at https://curatr.mcf.embl.de/. The library is freely available online and also integrated with other mass spectral repositories.

## INTRODUCTION

Metabolite identification remains a bottleneck in untargeted liquid chromatography-tandem mass spectrometry (LC-MS/MS) based untargeted metabolomics studies.^1^ This is one of the major challenges in converting untargeted mass spectrometry data into biological knowledge, inhibiting the growth of metabolomics compared to other “omics” sciences.^1,2^

Matching accurate mass (*m/z*), fragmentation (MS/MS) spectra and chromatographic retention time (RT) from a sample with authentic standards are considered as the gold standard Level 1 identification as formulated by the Metabolomics Standards Initiative.^2–4^ However, this approach is limited by the availability of authentic standards in individual labs. Alternately, achieving Level 2 of metabolite annotations by using standard reference mass spectral libraries is more attainable and thus plays a major role in LC-MS/MS-based untargeted metabolomics.^2,5^ Hence, public and commercial mass spectral libraries have become indispensable resources for fast and reliable compound identifications in untargeted metabolomics studies.^2,6^ Currently, a number of mass spectral libraries such as GNPS^7^, MoNA (includes MassBank^8^ and other databases), Metlin^9^, NIST-14^10^, mzCloud^11^, HMDB^12^, and LIPID MAPS^13^ are providing useful resources for metabolite and lipid identification.^2^

However, we found three major limitations while using available public libraries for untargeted metabolomic analysis. The first limitation is the lack of chromatographic information in most of the libraries. Most of the metabolomics studies use LC-MS/MS methods, however, many mass spectral libraries are generated from direct infusion MS/MS data thus lacking associated retention times or chromatographic profiles.^2^ Although the MoNA (MassBank) and NIST-14 (commercial) libraries have some chromatographic information included, none of these libraries provide visualization of chromatographic peaks of precursor ions (called mzRT features).^2^ This gap of knowledge is critical, as matching the chromatographic information such as RT from biological samples with standards acquired on similar LC-MS methods can improve the confidence in metabolite annotations.^14^ The chromatographic separation can also distinguish isobaric and isomeric metabolites which are indistinguishable by their accurate mass and even MS/MS spectra. Furthermore, the mobile phase buffers, solvents and additives can influence the formation of adduct ions and their fragmentation patterns.^6,15,16^ Another challenge in the LC-MS analysis is optimizing chromatographic retention, peak shape, and intensity which is often critical for reliable metabolite quantifications.^17–19^ Thus, LC-MS/MS library with chromatographic visualization for different adduct ions would assist in the method development for better annotation and deconvolution of untargeted metabolomic data as well as for assessing the quality of the individual LC-MS/MS data.^2,16^ Despite these potential benefits, visualization of the chromatographic profiles is not implemented in any of the public spectral libraries.

The second limitation is the lack of download access for some of the largest high-resolution MS/MS libraries such as METLIN and mzCloud which prevents using them both for batch search with in-house software and for developing new tools. Other libraries provide commercial access only, such as NIST-14, that limits the access compared to public spectral libraries.^2^

The third drawback is the presence of non-endogenous metabolites in the libraries which gives false positive and non-specific annotations for biological samples. Other than HMDB, most spectral libraries contain drugs, drug metabolites, synthetic organic compounds, toxicants, food additives, etc. which can have overlapping masses with endogenous metabolites.^20,21^

Here we report the first public LC-Orbitrap-MS/MS spectral library called EMBL-MCF (European Molecular Biology Laboratory-Metabolomics Core Facility). The library contains curated LC-MS/MS spectra of 435 endogenous metabolites and lipids. The unique feature of the library is its visualization of chromatographic peaks with their MS/MS spectra from different adducts. We used an open-source web application ‘*curatr*’ developed in-house to facilitate the creation, curation, and sharing of our spectral library.^22^ The library is freely available online and downloadable in multiple formats and is integrated into other repositories.

## EXPERIMENTAL

### Reagents, chemicals and metabolite standards

LC-MS grade water, methanol, isopropanol, and acetonitrile were purchased from BioSolve (Valkenswaard, The Netherlands). The MS grade mobile phase additives; ammonium formate, ammonium acetate, ammonium hydroxide, and formic acid were purchased from Sigma-Aldrich (Merck KGaA, Darmstadt, Germany). The list of all standards along with their vendor information is available online http://curatr.mcf.embl.de/inventory/. The Mass Spectrometry Metabolite Library of Standards (MSMLS) kit was purchased from IROA Technologies (MA, USA). All other metabolites and lipid standards were purchased from Sigma-Aldrich, Avanti Polar Lipids, and Matreya. HCT-116 (ATCC^®^ CCL-247) cells were culture to 80% confluency using vendor provided protocol (ATCC; VA, USA).

### Preparation of standards and samples

Multiplexed standard mixtures were prepared based on a chemical class of metabolites, suitability with LC-MS protocols, and non-overlapping molecular weights. About 1 mg/mL stock solutions were prepared in methanol/H2O (50:50; v/v) for primary metabolites and in methanol:isopropanol (50:50, v/v) for lipids. The equal volumes of stock solutions were combined to yield a mixture of 10 to 15 non-isobaric compounds at a concentration of about 10 μg/mL (standards mix). The ‘standards mix’ were then further 10x diluted in respective mobile phases to obtain a final concentration of about 1 μg/mL for each compound. In the case of IROA MSMLS kit; standards were diluted in water-methanol (50:50; v/v) or in methanol to get a final concentration of 25 μg/mL(as recommended by the supplier). The 10μL of these solutions were injected into the LC-MS system. In cases where mass spectrometry response was low, higher concentrations of metabolites were used. The metabolite extraction from ~2×10^6^ HC-T116 cells was performed as described in the earlier protocol.^23^

### LC-Orbitrap-MS/MS analysis

The library is created using six LC protocols in reversed-phase (RP) and hydrophilic interaction liquid chromatography (HILIC) separation modes for different classes of metabolites and lipids. The LC-MS parameters used for these protocols are summarized in Supplementary Methods. The data were acquired on Q-Exactive Plus mass spectrometer (Thermo Fisher Scientific, Bremen, Germany) coupled with either Infinity 1260 Series HPLC (Agilent Technologies, Palo Alto, CA) or Vanquish UHPLC system (Thermo Scientific, Massachusetts, USA). In most cases, the MS/MS spectra obtained from the top 10 most intense MS1 ions in the data-dependent acquisition (DDA) mode using inclusion lists. The stepped normalized high-energy dissociation (HCD) collision energies (CE) of 20, 40, 60 units were used. For each standard mixture data was acquired in both ESI positive and negative modes.

### Curating LC-MS/MS spectra

LC-MS/MS data was acquired using Thermo Xcalibur software (version 3.1, Thermo Electron Corporation). Our in-house developed web application ‘*curatr*’ was used for data curation, cataloging, and sharing the mass spectral library online.^22^

The overall workflow for creating the spectral library is described in Figure 1. All acquired LC-MS/MS raw files were converted to centroid mzML format^24^ using msConvert software (ProteoWizard version 3).^25^ A mzML file of standard mixtures was then submitted to the ‘*curatr*’ web-application.^22^ The metadata and other relevant parameters are listed in Supplementary Methods. The detailed video protocol for browsing, download, curation using ‘*curatr*’ web application is available on the website https://curatr.mcf.embl.de/.

**Figure 1.**
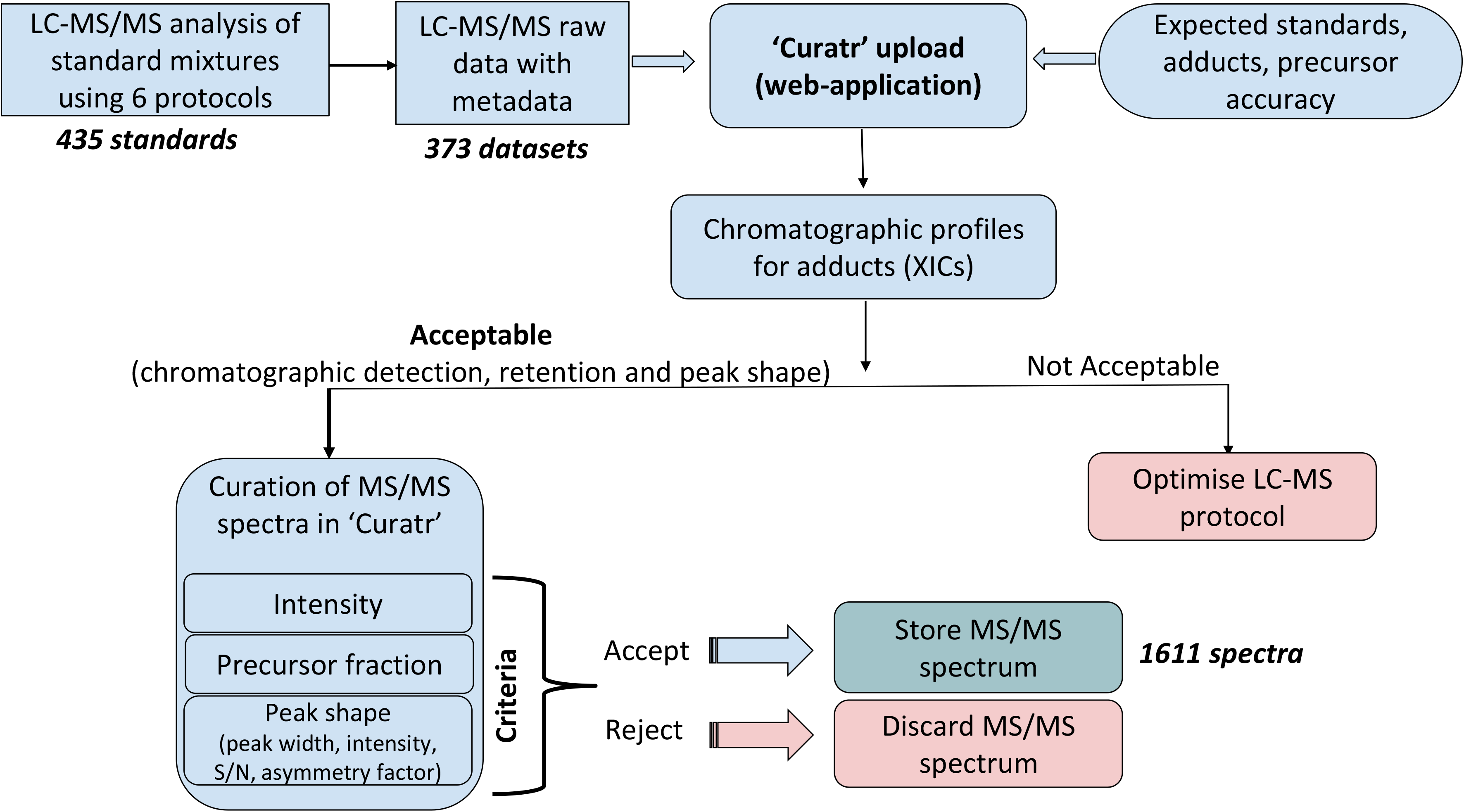
Workflow and criteria for curation of the EMBL-MCF LC-MS/MS spectral library by using the ‘curatr’ web application.

The curation of the LC-MS/MS spectra is a key part of this work and was performed by expert mass spectrometrists using online visualization and curation features in the ‘*curatr*’ web application. As shown in Figure 1, the first step of curation was an evaluation of the quality of extracted ion chromatogram (XIC) for optimal retention (>0.5 min), intensity (>105 units) and peak shape. We used three peak shape attributes namely; peak width or duration (>0.05 min), signal to noise (S/N >10) and peak asymmetry and/or tailing to evaluate chromatographic peak quality. Then MS/MS spectra collected in the data-dependent manner across the chromatographic peak were checked to find a MS/MS spectrum with the presence of precursor ion fraction (> 0.1). Next, each fragmentation spectra were interpreted manually based on their chemical structure and *in-silico* predictions using Mass Frontier Spectral Interpretation Software (version 7.0; HighChem Ltd., Bratislava, Slovakia). Some spectra were cross-checked with other public libraries, if available. Based on these criteria each MS/MS spectrum was either accepted or rejected. The accepted spectra were then stored with the molecular ID, InChI key, chromatographic information, MS parameters, PubChem CID, SPLASH spectral identifier, and solubility information.

### Visualizing LC-MS/MS spectra

The ‘*curatr*’ web application can be used not only for curating the spectra, but also enables access, visualization, querying, and download of the EMBL-MCF spectral library in a web browser. The library can be queried either by the molecular name, formula or PubChem CID and the results are output as a table, presenting the list of available spectra. For a particular molecule, multiple MS/MS spectra curated for the same standard using different LC-MS protocols which can be visualized systematically in a single web-browser window. For each curated MS/MS spectrum, the method details and overlay precursor XIC profiles are displayed in a web-browser window (Figure 2).

**Figure 2.**
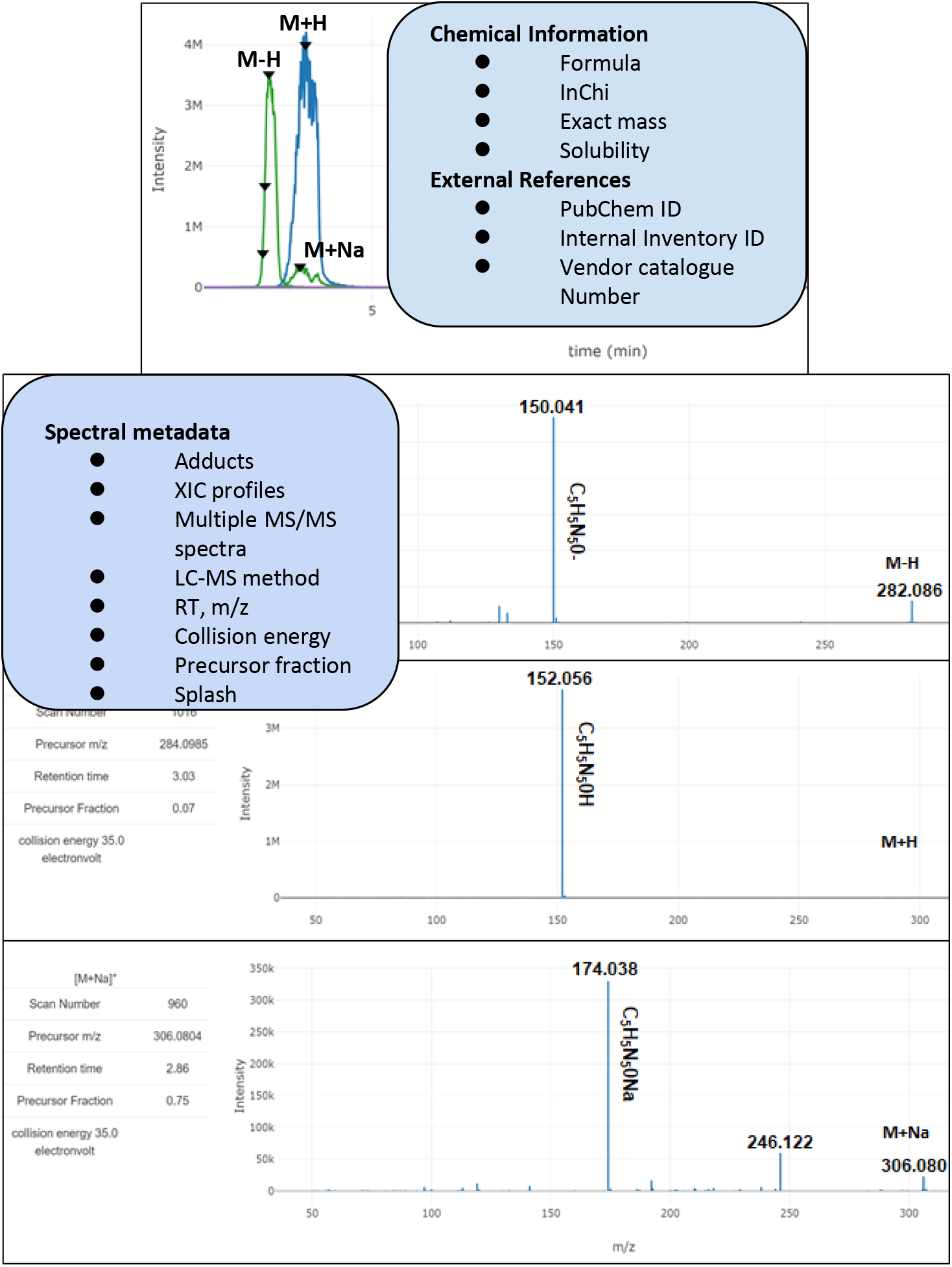
The ‘Standard detail’ layout in EMBL-MCF library for guanosine, as visualized using the ‘curatr’ web application. The upper panel shows an overlay of XICs of three ions of different adducts. The lower panel shows the MS/MS spectra for three detected adducts when using either negative or positive mode ([M-H]^-^, [M+H]^+^, and [M+Na]^+^). The triangles on XICs represent the data points where MS/MS spectra are acquired. The boxes display the type of metadata displayed with each library record.

## RESULTS AND DISCUSSION

The EMBL-MCF spectral library contains over 1600 MS/MS spectra from 435 authentic standards. These spectra were curated from 373 LC-MS/MS datasets acquired using six different LC-MS methods. The library has three unique features that distinguish it from other public mass spectral databases; 1) visualization of chromatographic peaks, 2) presence of different ion adducts, and 3) presence of LC-MS/MS data for unique 144 authentic standards which are not present in other open spectral libraries (MoNA and HMDB) (Supplementary Table 1).

The EMBL-MCF spectral library includes over 800 XICs of different ion adducts from 435 authentic standards. The library provides a visual reference for chromatographic peaks which can assist in assessing the quality of the data and assist in LC-MS method development. For example, Figure 2 shows an overlay of three adducts XICs ([M-H]^-^, [M+H]^+^, and [M+Na]^+^) of guanosine using two HILIC methods in the negative (with basic pH acetate buffer) and positive (with acidic pH formate buffer) ionization modes. For all three adducts, the exact *m/z* and MS/MS spectra are of consistently good quality, however, the XICs of positive mode adducts [M+H]^+^ and [M+Na]^+^ are noisy and of low intensity. This suggests that the [M-H]^-^ adduct obtained from the negative mode with a basic pH acetate buffer method is more suitable for reliable identification as well as quantification of guanosine. The other major advantage of LC analysis is the separation of isomers which are not distinguishable when using MS1 and MS/MS information alone (Supplementary Figure 1). Our library aims to provide a standard reference for chromatographic and MS/MS spectral information to assist LC-MS method development work along with metabolite identification.

The second feature of EMBL-MCF spectral library is the coverage of common adducts (Figure 3a) obtained from different LC-MS methods (Figure 3b). Adducts play an important role in the mzRT feature deconvolution, annotations, and even in structural identifications.^16^ Since most of the standards in the EMBL-MCF library were analyzed in both positive and negative ionization modes, we were able to detect six common adducts ([M-H]^-^, [M-H2O-H]^-^, [M+H]^+^, [M+Na]^+^, [M+NH4]^+^, [M+K]^+^). Figure 3a shows the ClassyFire^26^ based chemical class-specific distribution for six adducts detected from 435 standards. The deprotonated [M-H]^-^ adduct was the most common among organic acids, fatty acids, organic oxygen, and nucleotides, while [M-H2O-H]^-^ was found in a few organic oxygen and acid metabolites. The protonated adduct [M+H]^+^ was more common in lipids, heterocyclic, and nitrogen-containing organic metabolites. The ([M+Na]^+^, [M+K]^+^) adducts were mainly observed in lipids. The use of ammonium acetate and ammonium formate buffered mobile phases facilitated the formation of [M+NH4]^+^ adduct in both metabolite and lipid standards.

**Figure 3.**
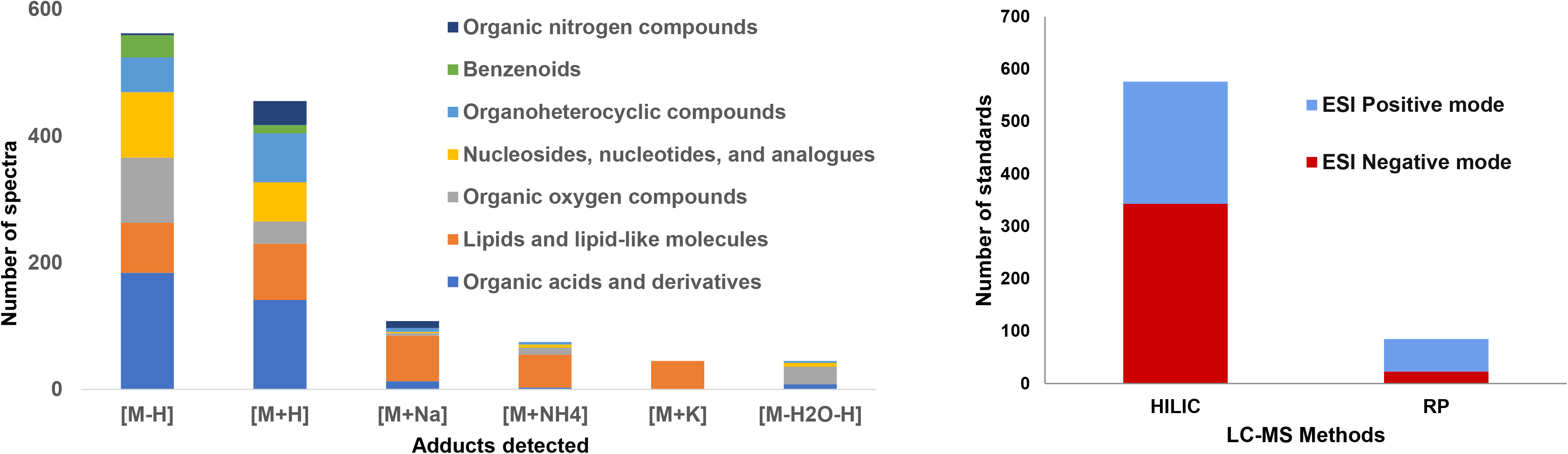
(a) ClassyFire based Chemical class-specific distribution of six common adducts detected from over 1600 MS/MS spectra obtained from 435 standards of the EMBL-MCF library. (b) The number of standards detected in both positive and negative ionization modes from the HILIC and RP based LC-MS methods used for the library generation (see Supplementary Methods for details).

The MS/MS data of different adducts from both positive and negative modes can play a complementary role in structural identifications. As shown in Figure 2, MS/MS data of three adducts [M+H]^+^, [M+Na]^+^, and [M-H]^-^ gives additional confirmation for the metabolite identification of guanosine. Also, in case of lipid identification, additional structural information for the assignment of fatty acyl chains can be obtained from different adducts. As shown in Figure 4, the structural elucidation of phosphocholine PC 16:0_18:1 using MS/MS spectra of two adducts [M+H]^+^ and [M+Na]^+^. The MS/MS spectrum of [M+H]^+^ confirms the presence of the head group (*i.e.,* phosphocholine at *m/z* 184.07) while MS/MS of [M+Na]^+^ supports the presence of fatty acyl chain C16:0. We have also previously illustrated the importance of having MS/MS reference data of different adducts from both ionization modes to enhance structural identification of lipids.^27^

**Figure 4.**
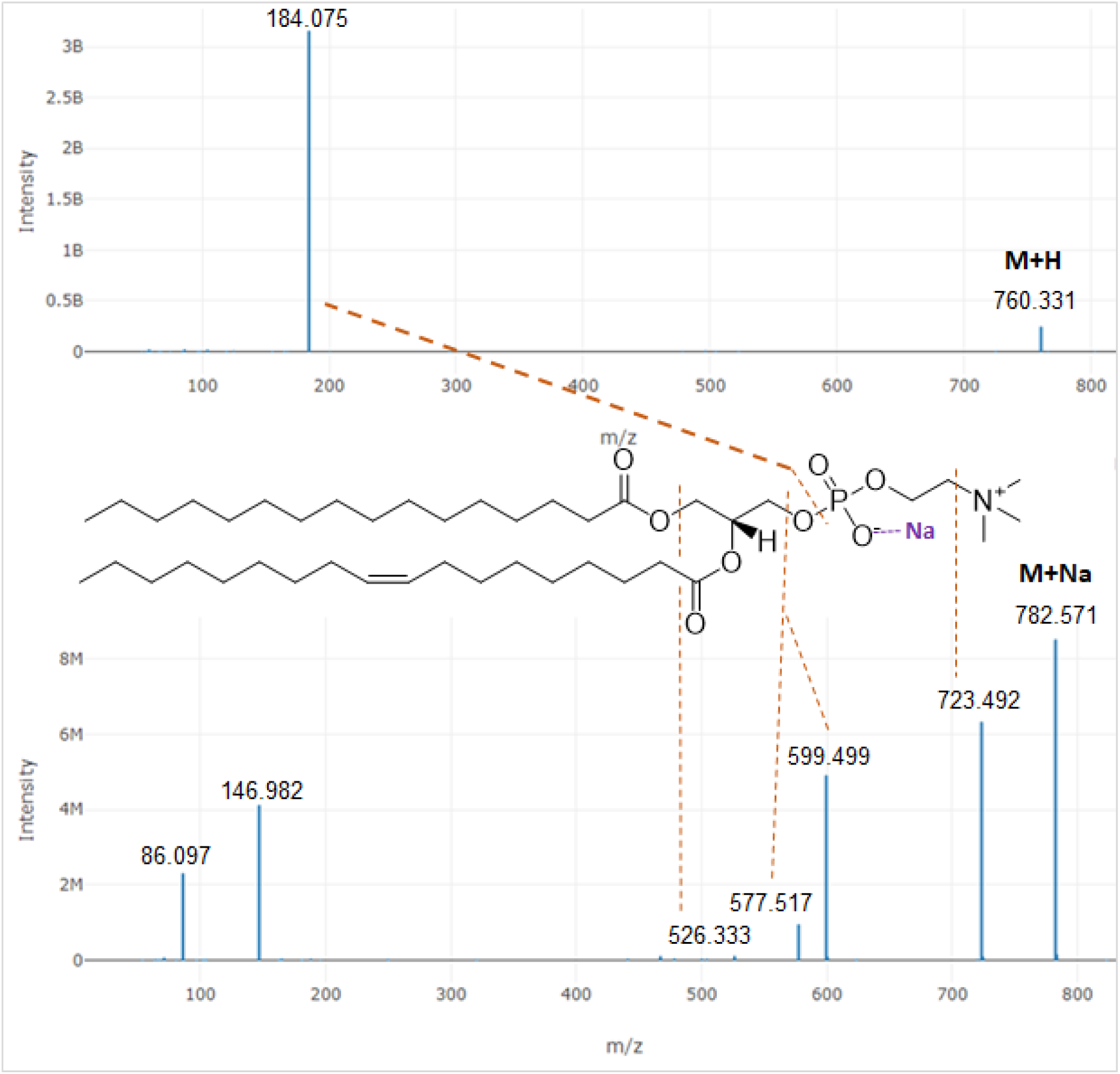
Complementary fragmentation spectra from two adducts ([M+H]^+^ and [M+Na]^+^) enhances the structural elucidation of a lipid (authentic standard of a phosphocholine PC 16:0_18:1).

The third important feature of the library is its focus on endogenous metabolites from biologically important chemical classes and pathways (Figure 5) with 37 unique metabolites present compared to other public libraries. The comparison was done with all the entries from MoNA database which is the collection of most of the open databases (Supplementary Information_Unique IDs).^2^ Although our library is much smaller compared to these two libraries, still it includes some unique metabolites and lipids which mainly represent lipids, organic acids, and nucleotides. It should be noted that HMDB, MoNa, and other databases do not provide visualization of chromatographic profiles (Supplementary Table 1). The selection of the metabolite standards in the library is performed in consultations with EMBL-MCF users who are mainly biologists and biochemists working in diverse fields of biology.^28^ The ClassyFire^26^ based chemical classification of the library metabolites shows broad chemical coverage (Figure 5) of the library. The library covers mainly carboxylic acids, nucleotides, glycerolipids, and fatty acids related to metabolites.

**Figure 5.**
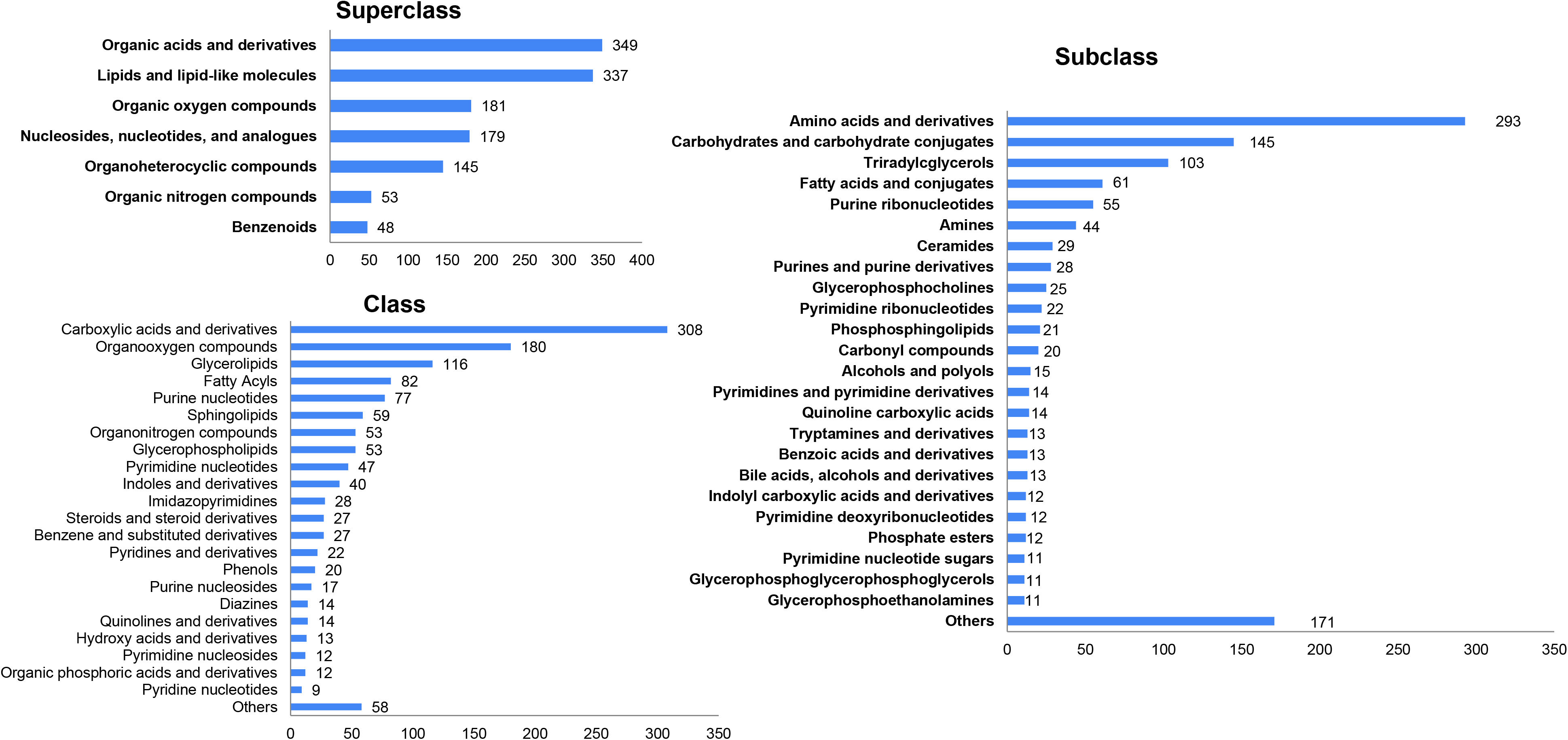
ClassyFire based classification of the EMBL-MCF spectral library showing chemical classes covered by the library.

### Chromatographic behavior of metabolites and lipids

The chemical diversity of metabolome poses a considerable challenge for chromatographic optimization in targeted and untargeted metabolomics analysis.^19^ Poor peak shapes significantly affect automated peak integration, data reliability, and increases the time required for manual peak curation.^17,19,29,30^ The selection of suitable chromatographic methods can reduce data analysis time and avoid unreliable results. However, the current method development approach relies on literature and expert trials. The general approach in metabolomics is the use of reversed-phase (RP) chromatography for lipids and secondary metabolites, and HILIC separation mode for the polar metabolites.^31^ We used six LC-MS methods for different classes of metabolites and lipids in both positive and negative ionization modes (Figure 3a). These methods can be divided into RP and HILIC with a combination of different buffers and column conditions (Figure 3b; Supplementary Methods).

We found that HILIC chromatography at basic pH (8.5-9.0) covers more number of metabolites from a single analysis as reported previously.^23^ We used HILIC mode with two types of stationary phases; amide (Waters BEH Amide column and amino (Phenomenex Luna Amino column). Although amino columns had better peak shapes and retention while the amide column was more stable over time and provided better sensitivity for some metabolites. The comparison of amide, amine, and zic-HILIC (Merck SeQuant^®^ ZIC-HILIC 3.5 μm; 100 × 2.1 mm) columns for amino acid analysis (Supplementary Figures 8-12) shows amide column is a good compromise between retention of polar metabolites, sensitivity, and analytical stability.

The lipid standards were chromatographically well behaved in RP mode with formate and acetate buffers for positive and negative ionization modes respectively. The fatty acids, in particular, showed a strong response in negative mode with ammonium acetate buffer compared to acidified formate buffer which is consistent with our previous report.^32^ Also, we have previously shown that complementary LC-MS/MS data in both positive and negative ionization modes is essential for the assignment of fatty acyl chains of some glycerophospholipids.^27^

### Web interface

The curation and visualization of 373 LC-MS/MS datasets acquired were one of the major challenges for the generation of this library. We used our in-house developed open-source web application ‘curatr’ to curate, store, and share the library.^22^ The web application assisted in manual curation of LC-MS/MS data and molecule-wise cataloging of all spectra from different analyses along with their metadata. The data can be browsed by metabolites or fragmentation spectra. A free-text query search can be performed for molecular formula or PubChem ID to retrieve the spectral information.

The web application shows data by using two layouts. The ‘Standard detail’ layout contains chemical information and external reference identifiers followed by an overlay of XIC profiles from all the datasets (Figure 2). The ‘Fragmentation Spectra’ layout includes spectral and LC-MS protocol details for displayed spectra (Supplementary Figure 3). Both layouts contain respective MS/MS spectra along with their precursor adducts and other MS raw data information. The ‘curatr’ web application is implemented using the Python Django library, is open-source, available on our GitHub repository, and can be installed externally or re-used.^22^

The library is freely available online https://curatr.mcf.embl.de/MS2/export/ to download in multiple data formats (.TSV, .MGF, .MSP). The positive and negative ion mode data can be downloaded separately and is recommended to be used this way to match the polarity used in the experiment. All the 373 LC-MS/MS raw data files along with method details are available at the MetaboLights repository (Study Identifier MTBLS1861). The library is also available on the MoNA and GNPS repositories.^7^

### Applications of the library

In the EMBL Metabolomics Core Facility, we use the library routinely for metabolite annotations from various biological samples in untargeted as well as targeted metabolomics studies.^27,28,33,34^ As an example, we analyzed the HCT-116 cell extract samples using our HILIC_BA2 protocol (Supplementary methods) and used our library for metabolite identification as well as eliminating false positive annotations. We used criteria of fragmentation match (score > 10%) of at least 1 fragment ion other than precursor ion (Supplementary Figures 3b-6b) and the RT match (with a tolerance of + 0.5 min). We showed several examples where false positive identifications were eliminated using RT match with the library standards (Supplementary Figures 3a-6a).

The library has also provided its utility within the GNPS framework to explore spectral similarity in LC-MS/MS data^35^ and for the creation of MS/MS reference spectra.^36^ Li *et al* in their ‘SubFragment-Matching method’ showed the use of the EMBL-MCF library as a benchmarking dataset for training machine learning models to identify metabolites in biological samples.^37^

Apart from metabolite identifications, the EMBL-MCF library and the ‘curatr’ software facilitate sharing spectra, selecting protocols, and maintaining an inventory of standards. For instance, users of our core facility often search and browse metabolites of their interest, whereas mass spectrometrists access protocols to decide on suitable LC-MS methods for their applications.^38^

### Limitations of the library

Here, we provide a well-curated LC-MS/MS library to facilitate confident metabolite annotations from datasets generated on high-resolving Orbitrap instruments using different LC-MS protocols. However, considerations should be given to the RT variations even when using similar LC protocols due to the inherent variability of chromatographic methods.^31^ Although the size of the library is comparatively small compared to other public libraries, it does include spectra for some unique metabolites (Supplementary Information_Unique IDs). We recommend using the library as a complementary resource with other public databases. Despite the library being developed exclusively on Orbitrap instruments, it can potentially complement MS/MS data generated from other mass spectrometry instruments such as Q-TOF. We used six LC-MS methods (Supplementary Methods) for different classes of metabolites and lipids in both positive and negative ionization modes (Figure 3). Further method development for chromatographically challenging metabolites is required using different combinations of buffers and column conditions. We were not able to optimize all the metabolites from IROA MSMSL kit due to either poor chromatography and/or low MS/MS spectral intensity of over 200 metabolite standards (Supplementary Information_IROA MSMLS kit).

## CONCLUSIONS

Here we report on a public high-resolution LC-MS/MS based ‘EMBL-MCF spectral library’. It is a highly curated Orbitrap library from 435 authentic standards covering broad chemical classes of metabolite and lipids. The library is multidimensional and includes LC-MS/MS data for common adducts acquired using different LC-MS protocols. This library provides visualization of chromatographic profiles along with MS/MS spectra which can assist in metabolite identification in general and specifically in assessing the quality of the MS/MS spectra and in developing LC-MS methods. The web-application *‘curatr’* was developed and used to assist in curation, storage, and sharing the library online. The availability of such open multidimensional LC-MS/MS data can be a useful resource for metabolomics researchers and computational scientists. For instance, this fragmentation data can be used to better understand the effect of LC-MS methods on fragmentation patterns. Such efforts can improve *in silico* RT prediction models and MS/MS-based structure elucidation to support high-confidence metabolite identification.

## ASSOCIATED CONTENT

## Supporting Information

The Supplementary Information data files, Supplementary Figures and Tables, and Supplementary Methods is available free of charge on the ACS Publications website.

## Data availability

The library can be freely downloaded in multiple formats at https://curatr.mcf.embl.de/MS2/export/. The raw data is available for download on Metabolights (Study ID: MTBLS1861) repository and individual spectral data can be downloaded from MoNA-MassBank of North America (https://mona.fiehnlab.ucdavis.edu/spectra/browse?query=tags.text%3D%3D%22EMBL-MCF%22&size=10) in .mgf or .msp formats. The code for our open-source spectral library curation tool is available on Github https://github.com/alexandrovteam/curatr

## ACKNOWLEDGMENTS

The content of the EMBL-MCF library is licensed under CC BY 4.0 by default. For more information, please visit the web page of the Creative Commons Attribution 4.0 International License (https://creativecommons.org/licenses/by/4.0/).

We thank Andreas Stefan Eisenbarth, Renat Nigmetzianov, and other members of the Alexandrov team (EMBL) for their software development and support, IT department (EMBL) for maintaining the operations of the server for the library. We thank members of the Gavin group, Schultz group, and Hentze group (EMBL) for providing some metabolite and lipid standards.

## Supplementary Methods

Supplementary Methods Table: LC-MS parameters for six protocols used for generating EMBL-MCF spectral library

Orbitrap Mass spec Parameters

Data analysis for online upload and curation of the LC-MS/MS datasets

## Supplementary Figures and Tables

Supplementary Table 1: Feature comparison of available spectral libraries with EMBL-MCF library.

Supplementary Figure 1: Separation of glucose-6-phosphate and fructose-6-phosphate isomers.

Supplementary Figure 2) Fragmentation Spectrum layout view

Supplementary Figure 3a) Identification of Inosine from HCT116 cell extract by matching exact mass and RT with standard XIC from the library

Supplementary Figure 3b) Identification of Inosine from HCT116 cell extract

Supplementary Figure 4a) Identification of Uridine from HCT116 cell extract by matching exact mass and RT with standard XIC from the library

Supplementary Figure 4b) Identification of Uridine from HCT116 cell extract

Supplementary Figure 5a) Identification of Glutamine from HCT116 cell extract by matching exact mass and RT with standard XIC from the library. The false positive peak at 8.70 min was rejected.

Supplementary Figure 5b) Identification of Glutamine from HCT116 cell extract

Supplementary Figure 6a) Identification of Asparagine from HCT116 cell extract by matching exact mass and RT with standard XIC from the library. The false positive peak at 9.4 min was rejected

Supplementary Figure 6b) Identification of Asparagine from HCT116 cell extract

Supplementary Figure 7) Curation of Malate peak: Although there is MS/MS match the annotation of Malate was rejected due to poor peak shape

Supplementary Table 2: List of metabolite analysed on amino, amide and zicHILIC columns for comparison of chromatographic performance.

Supplementary Figure 8) The comparison of amide, amine and zic-HILIC columns for amino acid analysis shows amide column as a good compromise between retention of polar metabolites, sensitivity, and analytical stability

Supplementary Figure 9) Comparison of chromatographic performance of 3 columns for analysis of Creatinine standard

Supplementary Figure 10) Comparison of chromatographic performance of 3 columns for analysis of Tryptophan standard

Supplementary Figure 11) Comparison of chromatographic performance of 3 columns for analysis of Carnosine standard

Supplementary Figure 12) Comparison of chromatographic performance of 3 columns for analysis of Glutamic acid standard

## Supplementary excel files

Supplementary Information_IROA MSMLS kit

Supplementary Information_Unique IDs

