## Supplementary Methods for "Public LC-Orbitrap-MS/MS Spectral Library for Metabolite Identification"

**Supplementary Methods Table:** LC-MS parameters for six protocols used for generating EMBL-MCF spectral library

| | | | | | | | | | |
|---|---|---|---|---|---|---|---|---|---|
| **Mobile Phase buffers** | Buffer A | 10 mM ammonium acetate in water | 0.1% Formic acid in Water | 10 mM Ammonium acetate+0.1% NH4OH (pH 9) | 10 mM ammonium acetate (pH4 with acetic acid) in water | 10mM Ammonium formate+0.2% Formic acid in (50:50) ACN: Water | 10mM Ammonium formate+0.1 % formic acid in Isopropanol: ACN (90:10) | 10mM Ammonium acetate in Isopropanol: ACN (09:10) | **ESI mode** |
| | Buffer B | 10 mM ammonium acetate in ACN | 0.1% Formic acid in ACN | ACN | 0.1 % acetic acid in Methanol | 10 mM Ammonium formate+0.2% Formic acid in (95:5) ACN: Water | 10mM Ammonium formate+0.1 % formic acid in Water:ACN (60:40) | 10mM Ammonium acetate in Water:ACN (60:40) | |
| **Column** | Ascentis C18 (10cmx 2.1mm; 2.7uM) | **RP_NA** | | | | | | **LIP_NI** | Negative |
| | Kinetex C18 (100x 2.1mm; 2.6uM) | | **RP_AA** | | **RP_AM** | | **LIP_AI** | | Positive |
| | Xbridge Amide (100x 2.1mm; 2.6uM) | | | **HILIC_BA1** | | | | | Negative/ Positive |
| | Kinetex HILIC (100x 2.1mm; 2.6uM) | | | | | **HILIC_AA** | | | Positive |
| | Luna Amino (150x 2 mm; 3uM) | | | **HILIC-BA2** | | | | | Negative/ Positive |

**Other LC parameters:**

**RP_AA:**

Column Temperature: 30℃

Flow: 0.3 ml/min

Gradients:

| Time | % Buffer B |
|---|---|
| 0 | 95 |
| 3 | 95 |
| 18 | 25 |
| 23 | 2 |
| 30 | 2 |
| 31 | 95 |
| 35 | 95 |

**RP_NA:**

Column Temperature: 30℃

Flow: 0.26 ml/min

Gradients:

| Time | % Buffer B |
|---|---|
| 0 | 95 |
| 3 | 95 |
| 18 | 25 |
| 23 | 2 |
| 30 | 2 |
| 31 | 95 |

| | |
|---|---|
| 35 | 95 |

**RP03:**

Column Temperature: 30℃

Flow: 0.3 ml/min

Gradients:

| Time | % Buffer B |
|---|---|
| 0 | 5 |
| 1 | 5 |
| 7 | 54 |
| 8 | 90 |
| 11 | 90 |
| 11.5 | 5 |
| 14 | 5 |

**RP04:**

Column Temperature: 40℃

Flow: 0.26 ml/min

Gradients:

| Time | % Buffer B |
|---|---|
| 0 | 5 |
| 3 | 5 |
| 10 | 75 |

| | |
|---|---|
| 12 | 98 |
| 16 | 98 |
| 16.1 | 5 |
| 19 | 5 |

**HILIC_BA:**

Column Temperature: 30℃

Flow: 0.26 ml/min

Gradients:

| Time | % Buffer B |
|---|---|
| 0 | 85 |
| 2 | 85 |
| 12 | 10 |
| 14 | 10 |
| 14.1 | 85 |
| 16 | 85 |

**HILIC_AA:**

Column Temperature: 30℃

Flow: 0.2 ml/min

Gradients:

| Time | % Buffer B |
|---|---|
| 0 | 100 |
| 5 | 100 |
| 11 | 30 |
| 14 | 30 |
| 15 | 100 |
| 18 | 100 |

**LIP_AI:**

Column Temperature: 30℃

Flow: 0.26 ml/min

Gradients:

| Time | % Buffer B |
|---|---|
| 0 | 20 |
| 0.5 | 20 |
| 3 | 50 |
| 10 | 70 |
| 18 | 97 |
| 22 | 97 |
| 22.1 | 20 |

| | |
|---|---|
| 25 | 20 |

**LIP_NI:**

Column Temperature: 30℃

Flow: 0.26 ml/min

Gradients:

| Time | % Buffer B |
|---|---|
| 0 | 32 |
| 1.5 | 32 |
| 4 | 45 |
| 5 | 52 |
| 8 | 58 |
| 11 | 66 |
| 14 | 70 |
| 18 | 75 |
| 21 | 97 |
| 25 | 97 |
| 25.1 | 32 |
| 28 | 32 |

**Q-exactive Orbitrap Mass spec Parameters**

Molecules were detected with HRMS full scan at the mass resolving power R=70000 in the mass range of 60-900 m/z (for small molecules) or 150 to 1500 m/z (for lipids). The data-dependent (DDA) MS/MS scans for 10 most intense (TOP10)were obtained along with full scans using higher-energy collisional dissociation (HCD) of normalized collision energies (NCE) of 20, 40, and 60 units at the mass resolving power R=17500. The MS parameters in the Tune software (Thermo Scientific) were set as; spray voltage of 4.2 kV (for negative mode 3.5 kV), sheath gas 35, and auxiliary gas 5 units, S-Lens 65 eV, capillary temperature 350$^{o}$C, and vaporization temperature of auxiliary gas was set at 300$^{o}$C.

**Data analysis for online upload and curation of the LC-MS/MS datasets**

All acquired LC-MS/MS .raw files were converted to centroid .mzml format using Proteowizard software. A .mzml file for standard mixtures was then uploaded on curatr web-application and respective standards were selected from curatr database inventory (http://curatr.mcf.embl.de/inventory/). The other parameters and metadata relevant to the data file were selected as:

1. Name of the standard (or MCF ID)
2. Expected adduct (based on LC-MS method used)
3. Anticipated MS accuracy for precursor (default value 10 ppm)
4. Precursor quadrupole window (default value 1 Da)
5. LC method information
6. MS method information
7. ESI ionization mode
8. MS ion analyzer used (default value QFT)

The parameters 1 to 4 are used by curatr software for extracting expected precursors, their adducts, and MS/MS spectra from a dataset. The input information 4 to 8 is stored with each dataset and spectra as LC-MS metadata.
