## Supplementary Figures and Tables for "Public LC-Orbitrap-MS/MS Spectral Library for Metabolite Identification"

**For sections ‘Chromatographic behavior of metabolites and lipids’ and ‘Applications of the library’**

Supplementary Table 1: Feature comparison for spectral libraries

| Library | LC info | Open access | Orbitrap |
|---|---|---|---|
| mzCloud | ✗ | – | ✔ |
| NIST14 | ✔ | ✗ | – |
| Metlin | ✗ | – | ✗ |
| Massbank/MoNa | – | ✔ | – |
| GNPS | ✗ | ✔ | – |
| HMDB | ✗ | ✔ | ✗ |
| EMSL (this library) | ✔ | ✔ | ✔ |

Supplementary Figure 1: Separation of isomers Glucose 6-phosphate and Fructose 6-phosphate

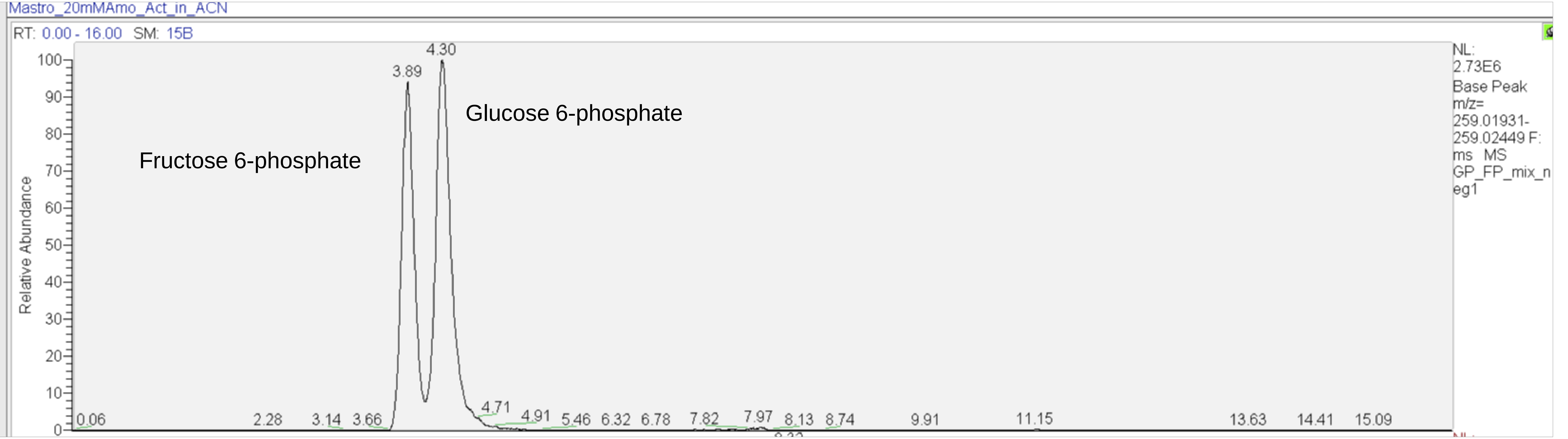


Column: C18 Shimadzu Mastro column
Mobile phase-A: 20mM ammonium acetate
Mobile phase-B:  Acetonitrile
ESI negative mode
Other parameters are same as RP_NA method (Supplementary Table 1)

Supplementary Figure 2) Fragmentation Spectrum layout view

Fragmentation Spectrum for 44: Guanosine, [M-H]-

Scan number: 1277

Precursor m/z: 282.08420

Retention time: 3.38 min

Collision energy: collision energy 30.0 electronvolt

Raw data: Std_ADP_phos_mix_170117112458.mzML

Splash: calculating...

LC Method: Column: Xbridge Amide (100X 2.1mm; 2.6uM); Temperature: 30 C; Mobile phase A: 7.5 mM Ammonium acetate+0.1% NH4OH, Mobile phase B: ACN; Flow: 0.3 ml/min Gradient : (Time/% B : 0/ 85, 2/ 85, 12/ 10, 14/ 10 , 14.1/ 85, 16/ 85)

MS Method: ESI_Neg

Review Information

Precursor intensity: 267338

Precursor Fraction: 1.0

Date added: April 27, 2017

Date curated: April 28, 2017

Curator: prasad

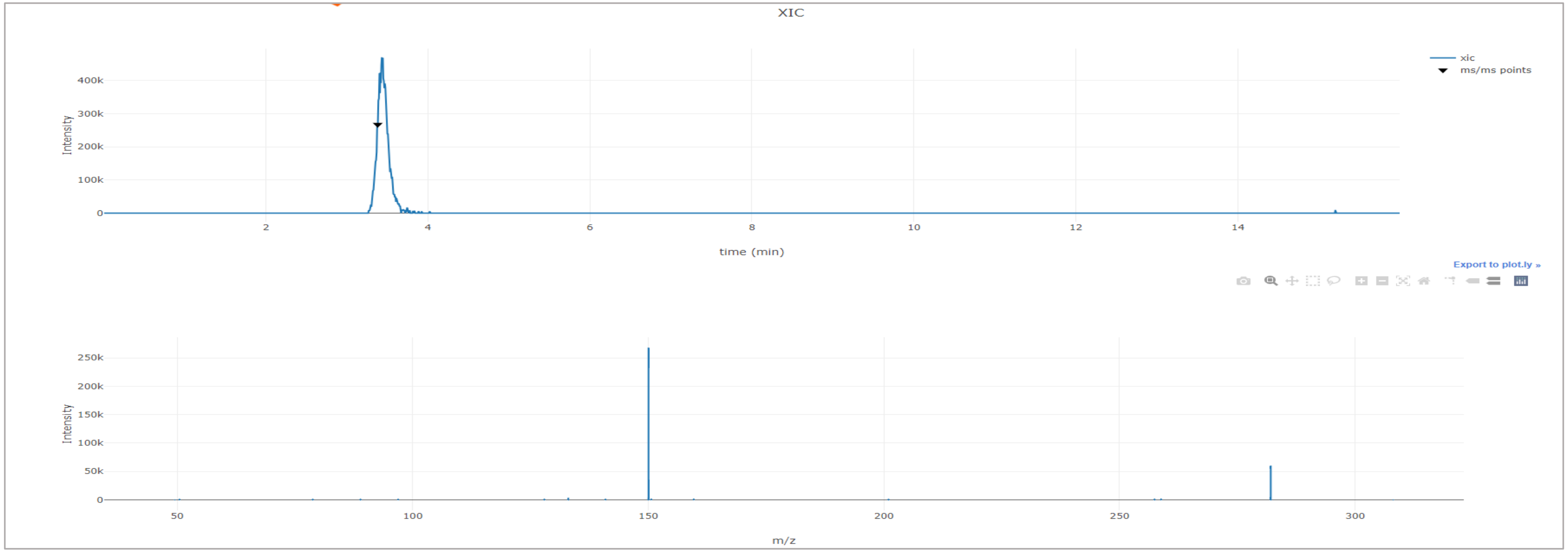

Supplementary Figure 3a) Identification of Inosine from HCT116 cell extract by matching exact mass and RT with standard XIC from the library

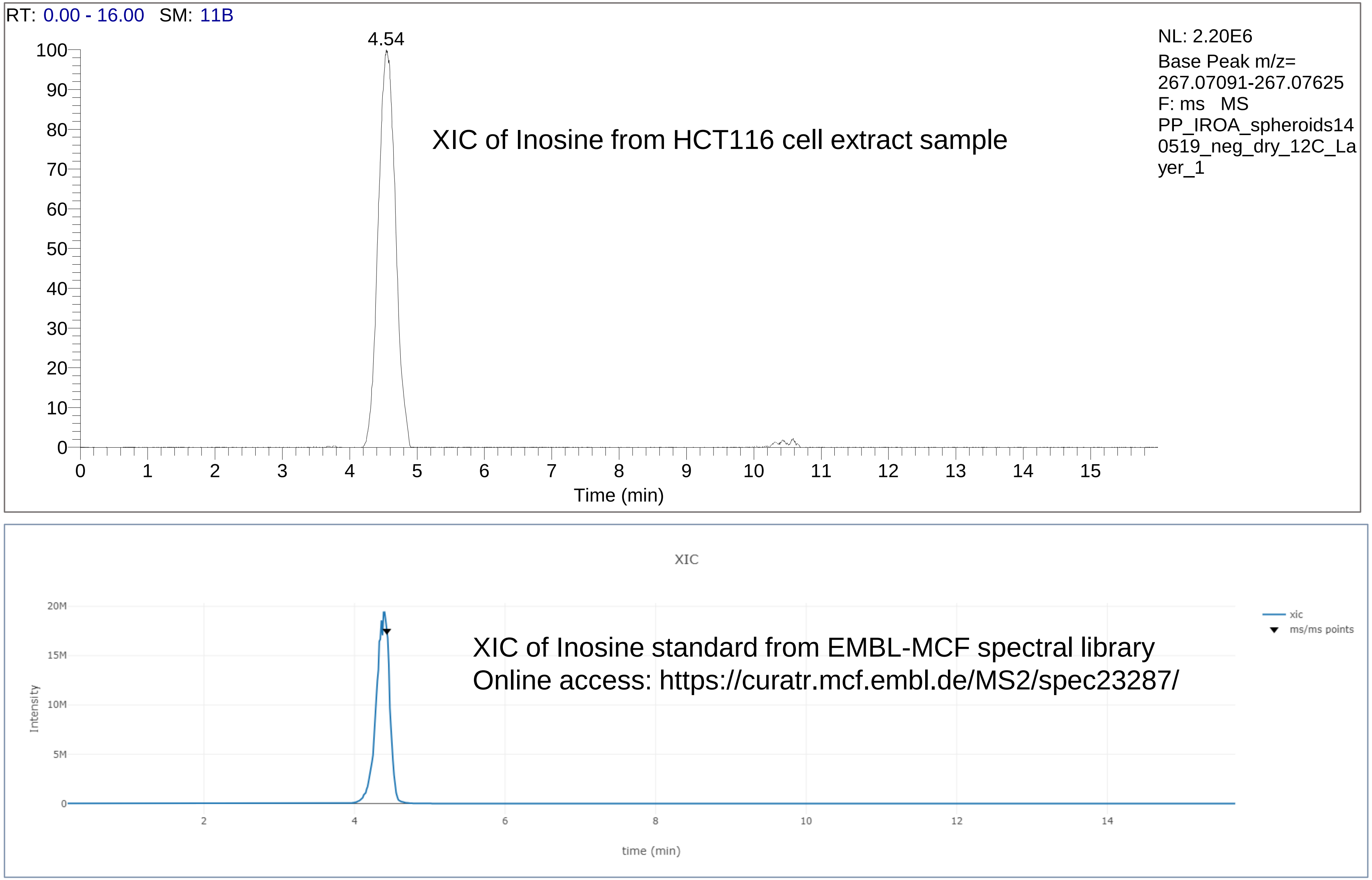

Supplementary Figure 3b) Identification of Inosine from HCT116 cell extract

**Compound 4.54_267.0729m/z**

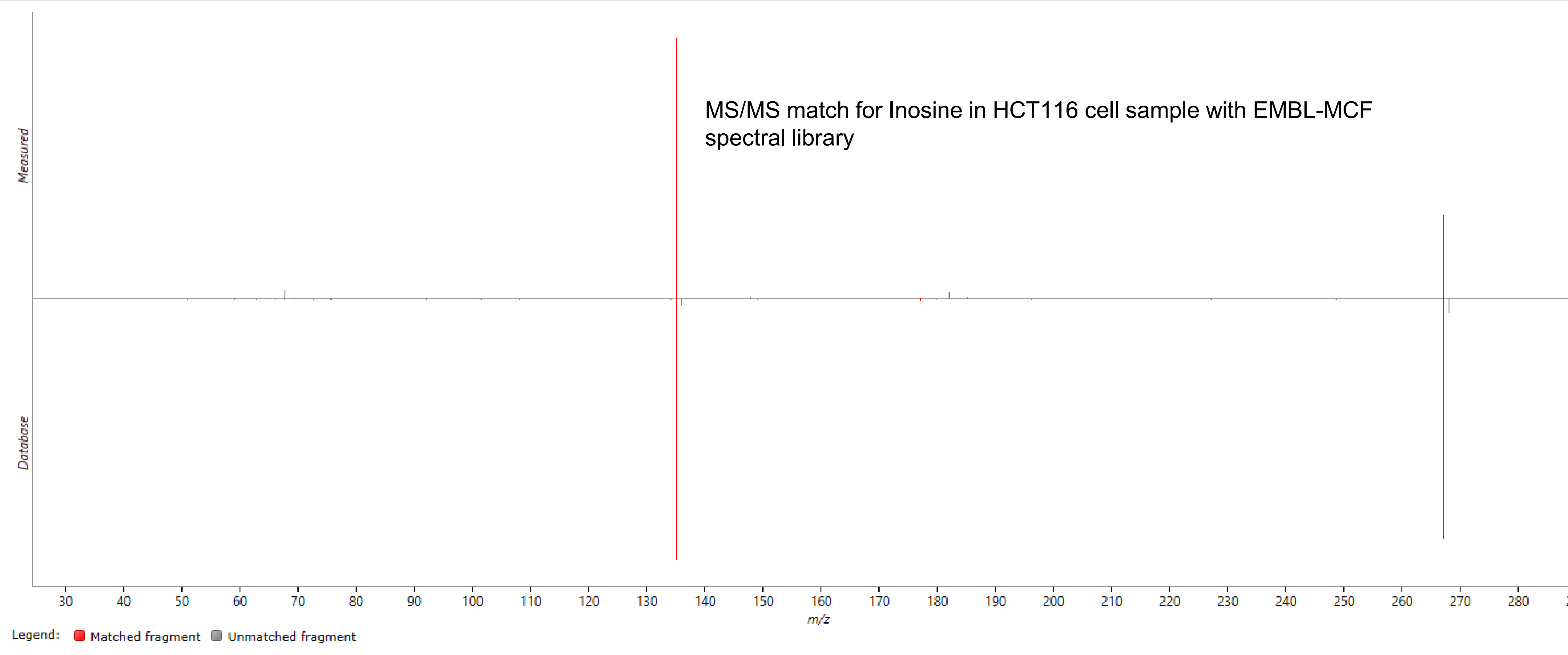


**Possible identifications: 1**

| ☆ | Compound ID | Description | Adducts | Formula | Retention time | Score | Fragmentation score | Mass error (ppm) | Re | Isotope similar | Link | Search Configuration |
|---|---|---|---|---|---|---|---|---|---|---|---|---|
| ★ | INOSINE | INOSINE | M-H | $C_{10}H_{12}N_4O_5$ | | 52.5 | 69 | -2.21 | | 95.93 | | Method: Progenesis MetaScope. Database: EMBL_MCF_neg.msp. Precursor tolerance: 10 ppm. |

Progenesis QI software screenshot with Fragmentation scores

Supplementary Figure 4a) Identification of Uridine from HCT116 cell extract by matching exact mass and RT with standard XIC from the library

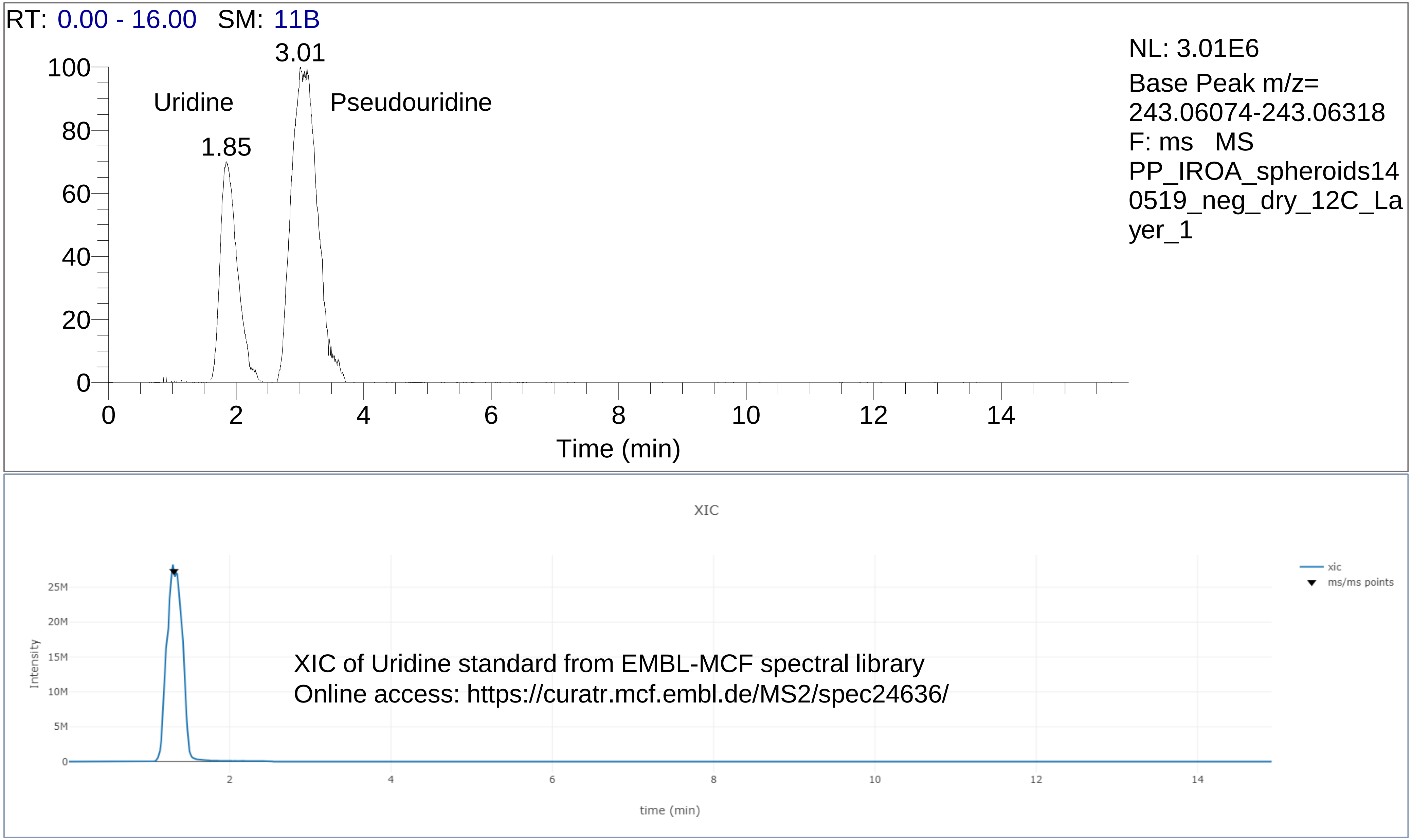

Supplementary Figure 4b) Identification of Uridine from HCT116 cell extract

**Compound 1.86_243.0616m/z**

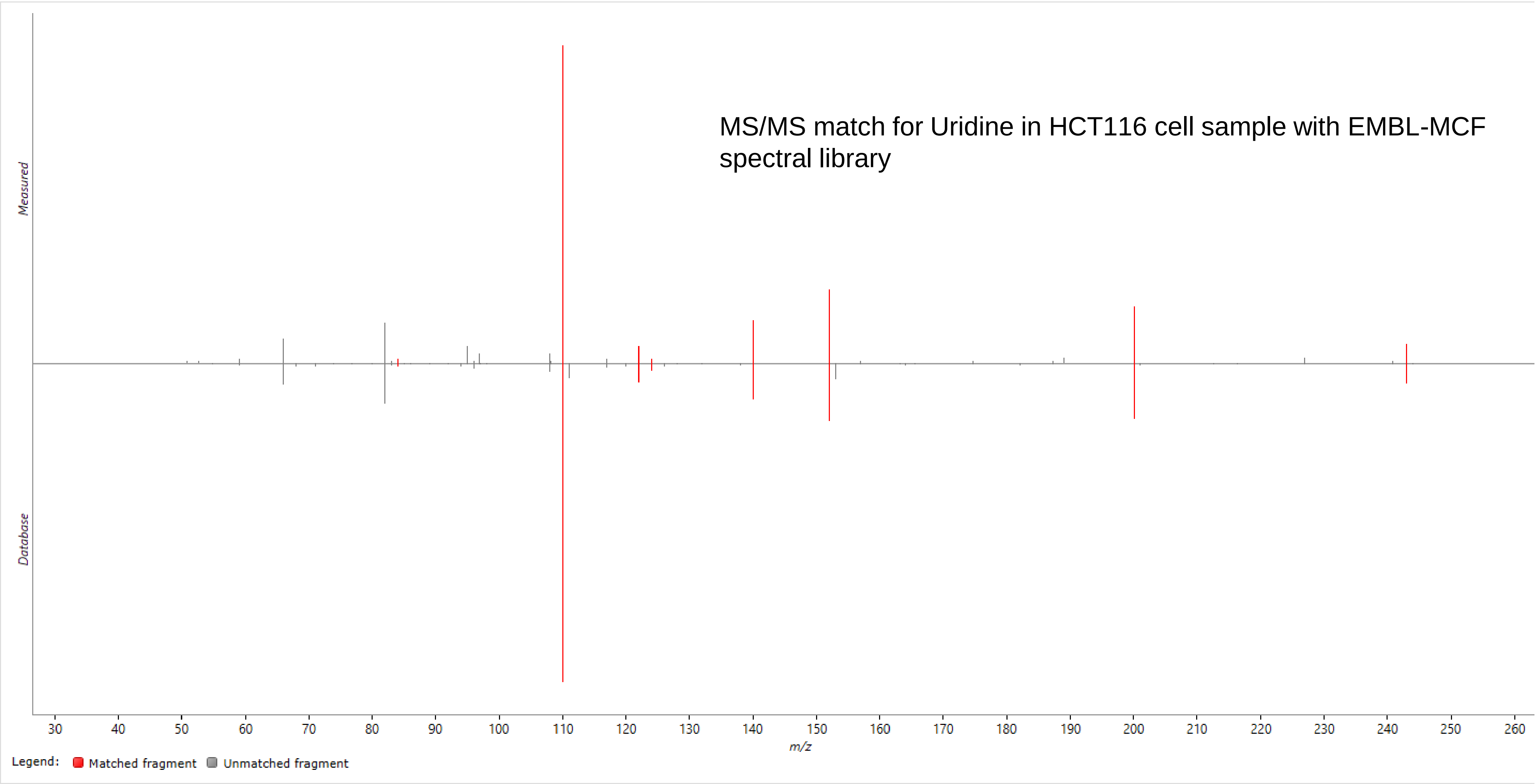


**Possible identifications: 1**

| ☆ | Compound ID | Description | Adducts | Formula | Retention time | Score | Fragmentation score | Mass error (ppm) | Re | Isotope similar | Link | Search Configuration |
|---|---|---|---|---|---|---|---|---|---|---|---|---|
| ★ | URIDINE | URIDINE | M-H | $C_9H_{12}N_2O_6$ | | 55.7 | 83.9 | -2.72 | | 97.63 | | Method: Progenesis MetaScope. Database: EMBL_MCF_neg.msp. Precursor tolerance: 10 ppm. |

Progenesis QI software screenshot with Fragmentation scores

Supplementary Figure 5a) Identification of Glutamine from HCT116 cell extract by matching exact mass and RT with standard XIC from the library. The false positive peak at 8.70 min was rejected.

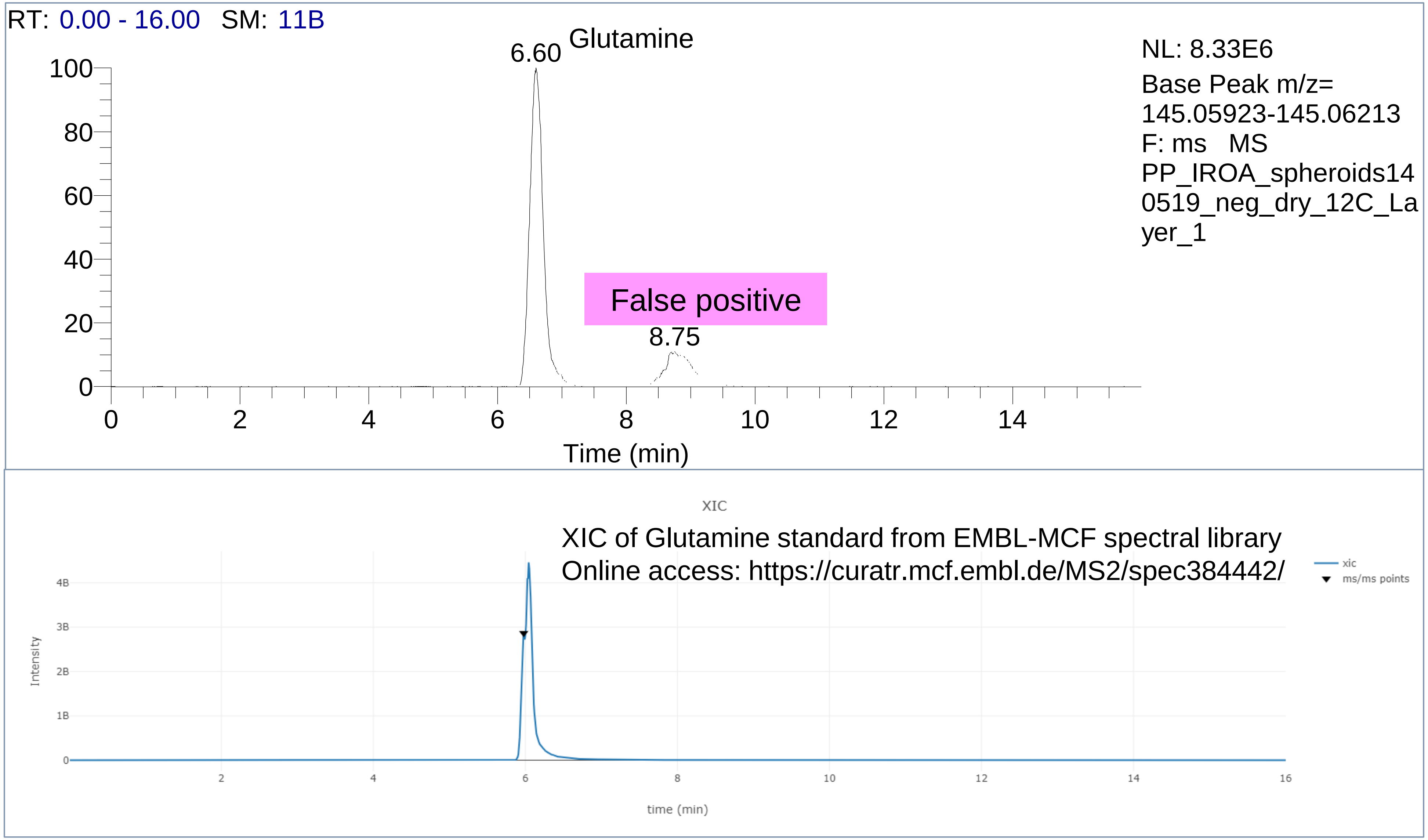

Supplementary Figure 5b) Identification of Glutamine from HCT116 cell extract

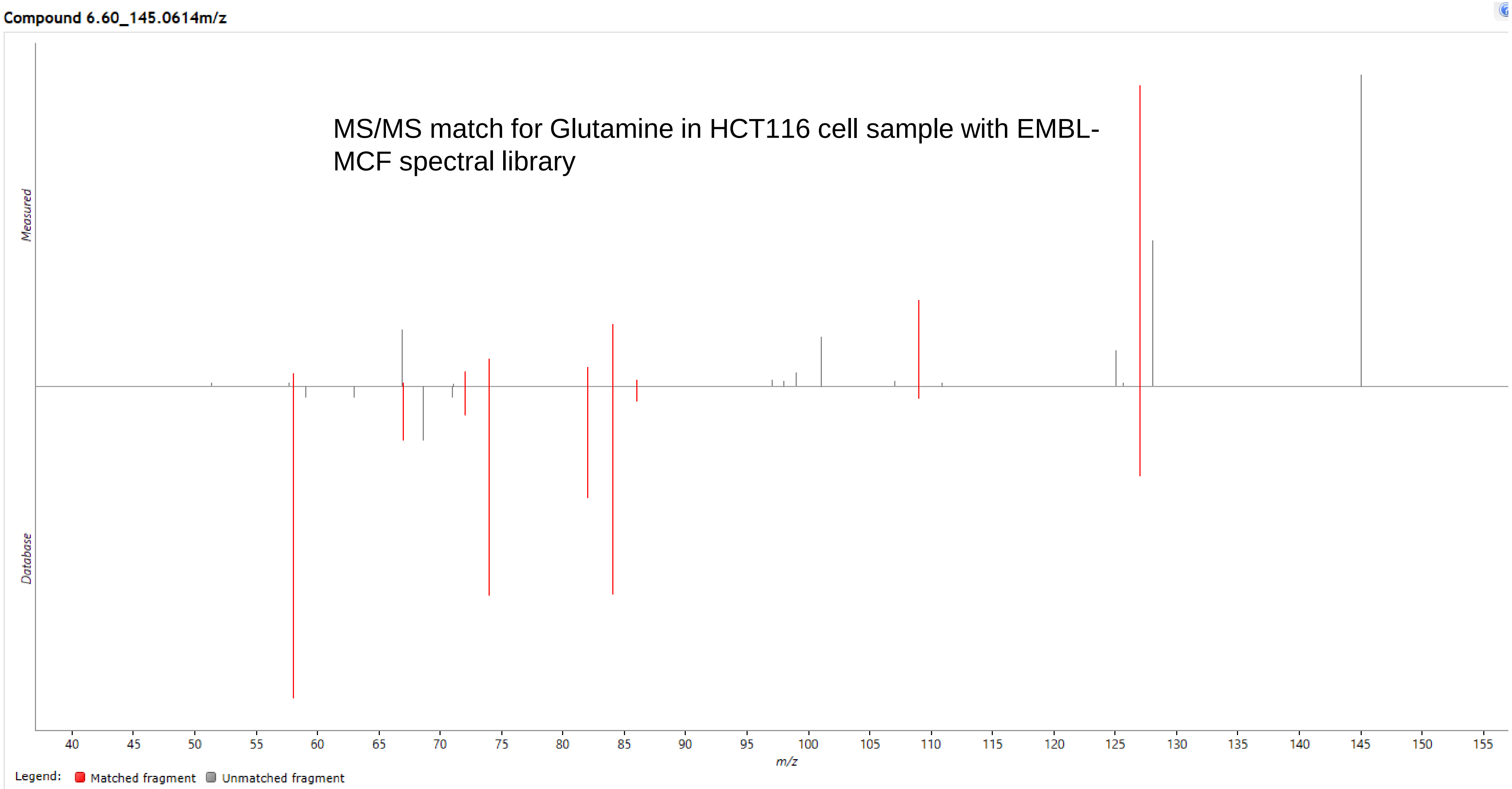


**Possible identifications: 1**

| ☆ | Compound ID | Description | Adducts | Formula | Retention time | Score | Fragmentation score | Mass error (ppm) | Re | Isotope similar | Linl | Search Configuration |
|---|---|---|---|---|---|---|---|---|---|---|---|---|
| ★ | L-GLUTAMINE | L-GLUTAMINE | M-H | $C_5H_{10}N_2O_3$ | | C 51.9 | C 64.6 | -2.96 | | 98.23 | | C Method: Progenesis MetaScope. Database: EMBL_MCF_neg.msp. Precursor tolerance: 10 ppm |

Progenesis QI software screenshot with Fragmentation scores

Supplementary Figure 6a) Identification of Asparagine from HCT116 cell extract by matching exact mass and RT with standard XIC from the library. The false positive peak at 9.4 min was rejected.

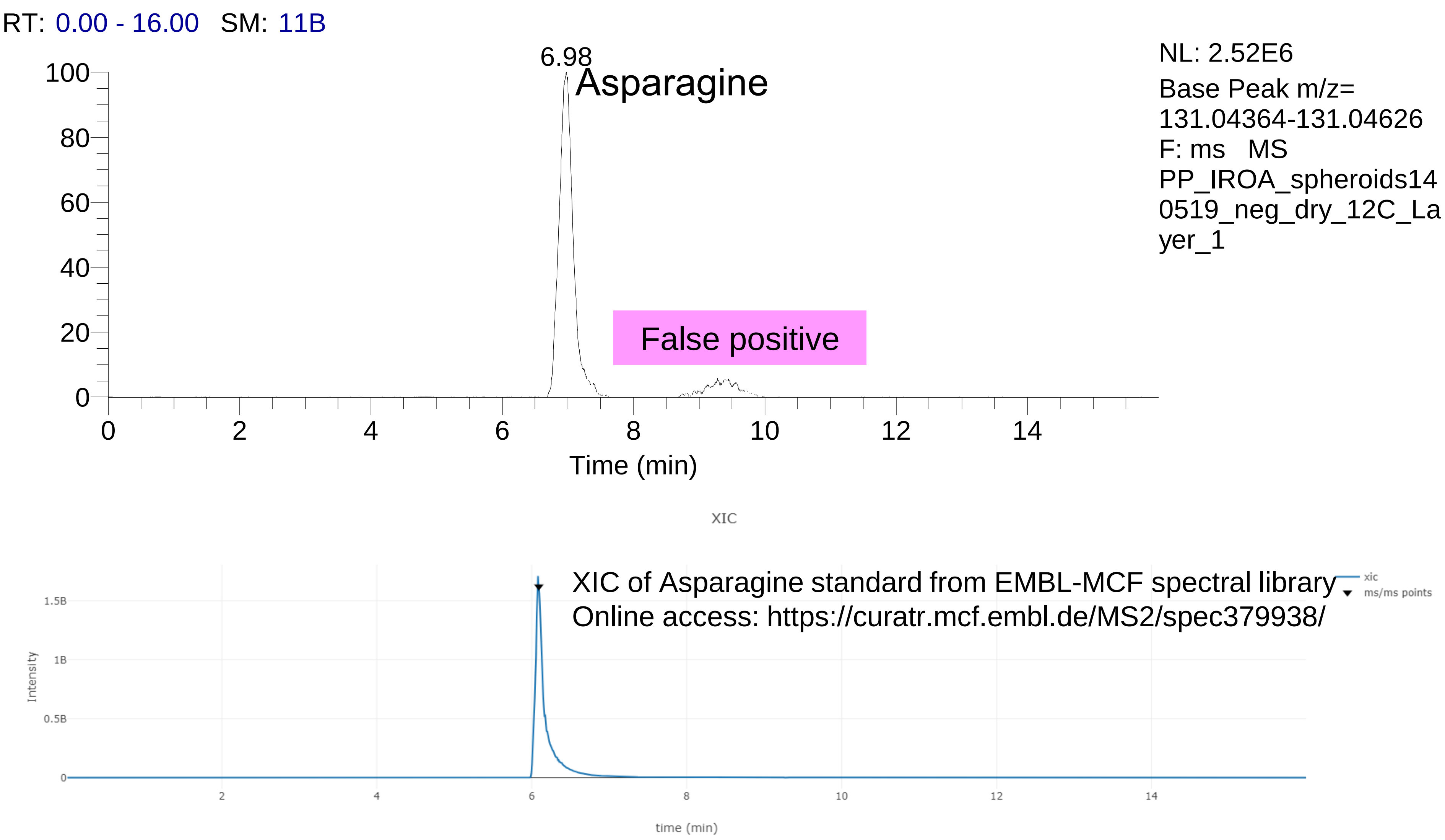

Supplementary Figure 6b) Identification of Asparagine from HCT116 cell extract

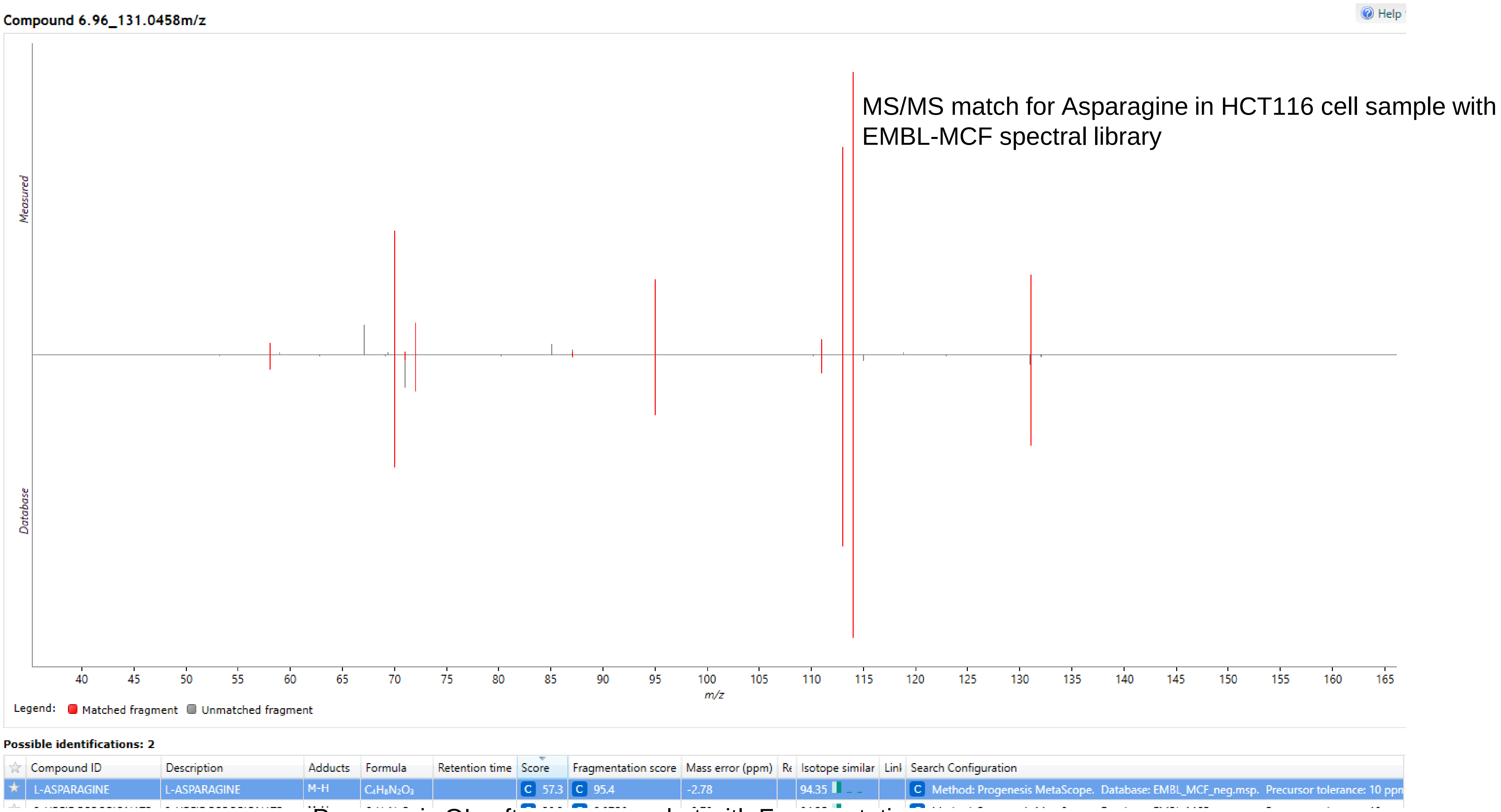


Progenesis QI software screenshot with Fragmentation scores

Supplementary Figure 7) Curation of Malate peak: Although  there is MS/MS match the annotation of Malate was rejected due to poor peak shape.

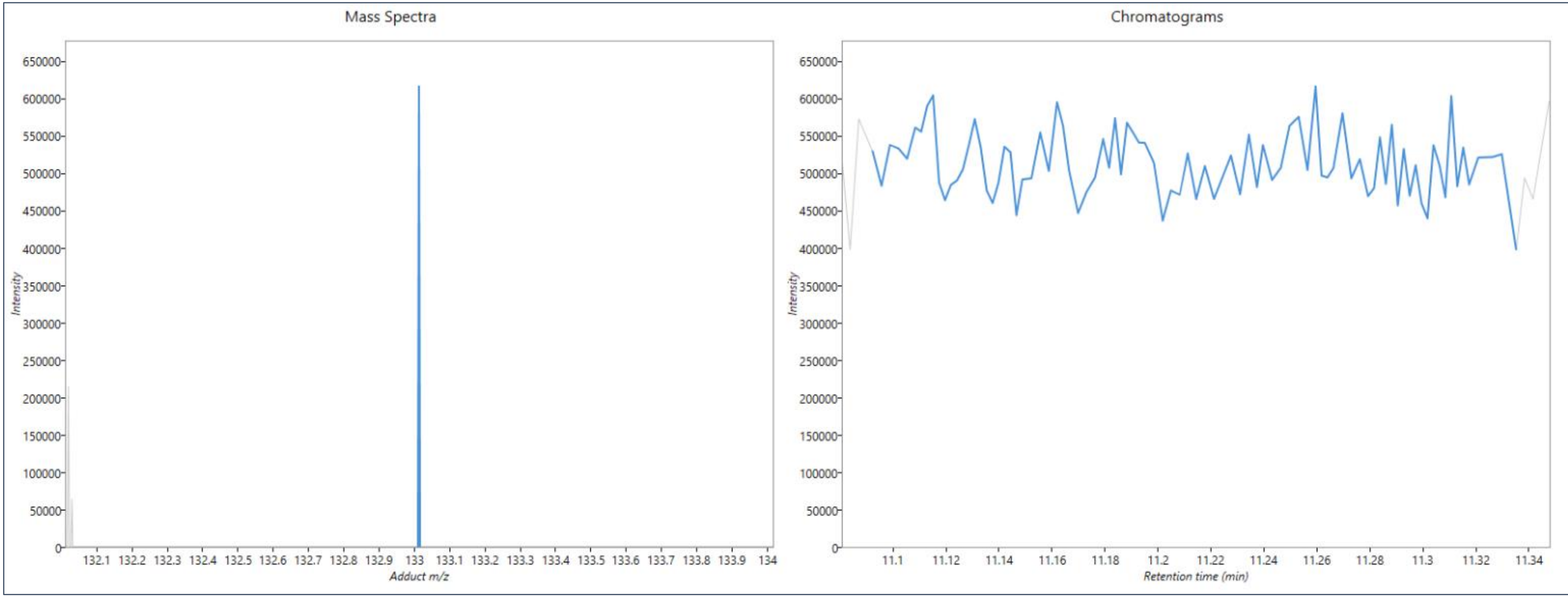

Supplementary Table 2: List of metabolite analysed on amino, amide and zicHILIC columns for comparison of chromatographic performance.

1. Glycine
2. Alanine
3. Sarcosine
4. Amino-n-butyric acid
5. Serine
6. Creatinine
7. Proline
8. Valine
9. Threonine
10. Taurine
11. Hydroxy-proline
12. Leucine
13. Ornithine
14. Aspartic acid
15. Homocysteine
16. Lysine
17. Glutamic acid
18. Methionine
19. Histidine
20. Aminoadipic
21. Hydroxylysine
22. Phenylalanine
23. Methyl histidine
24. Arginine
25. Citrulline
26. Tyrosine
27. Tryptophan
28. Cystathionine
29. Carnosine
30. Cystine
31. Anserine

Supplementary Figure 8) The comparison of amide, amine and zic-HILIC columns for amino acid analysis shows amide column as a good compromise between retention of polar metabolites, sensitivity, and analytical stability

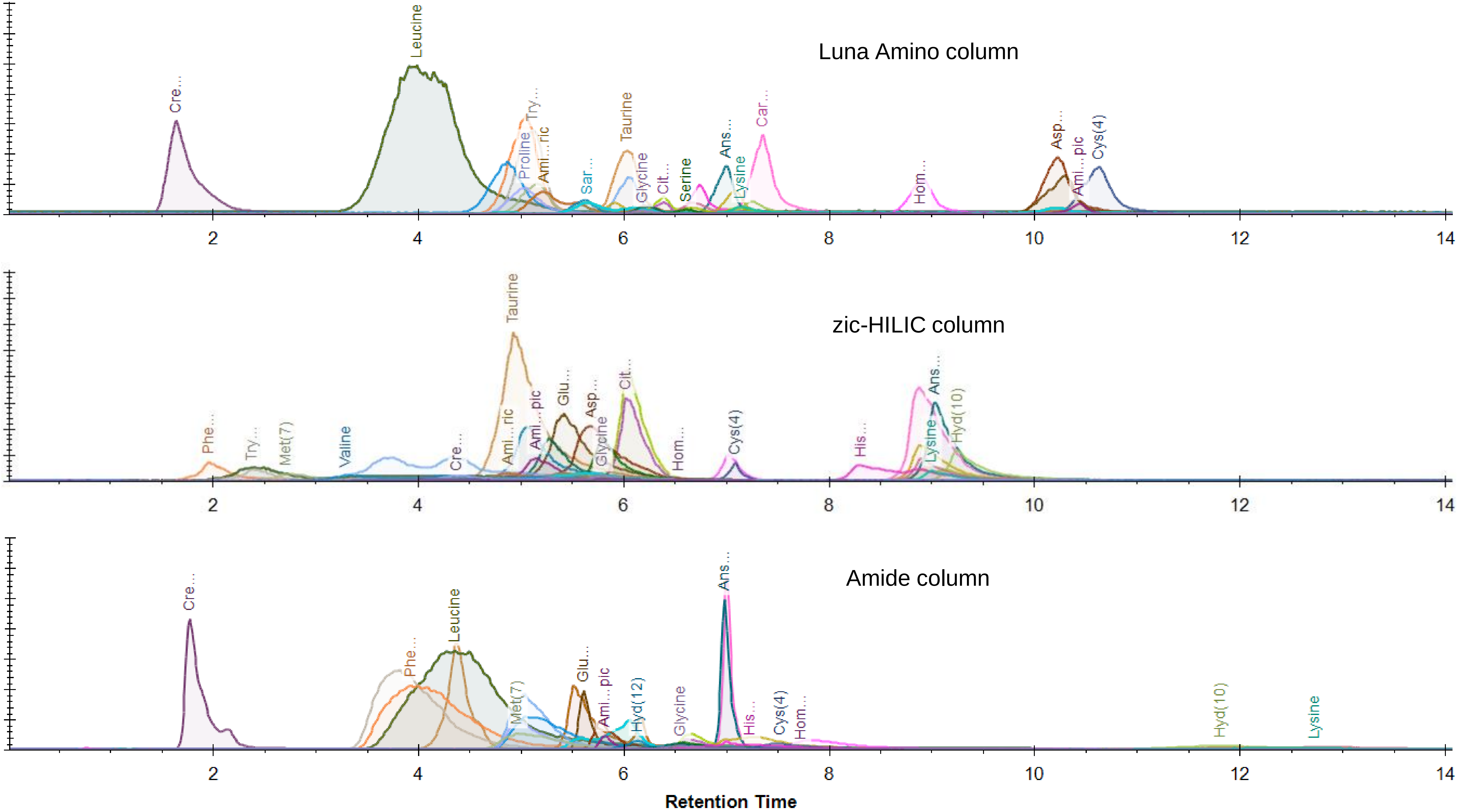

Supplementary Figure 9) Comparison of chromatographic performance of 3 columns for analysis of Creatinine standard

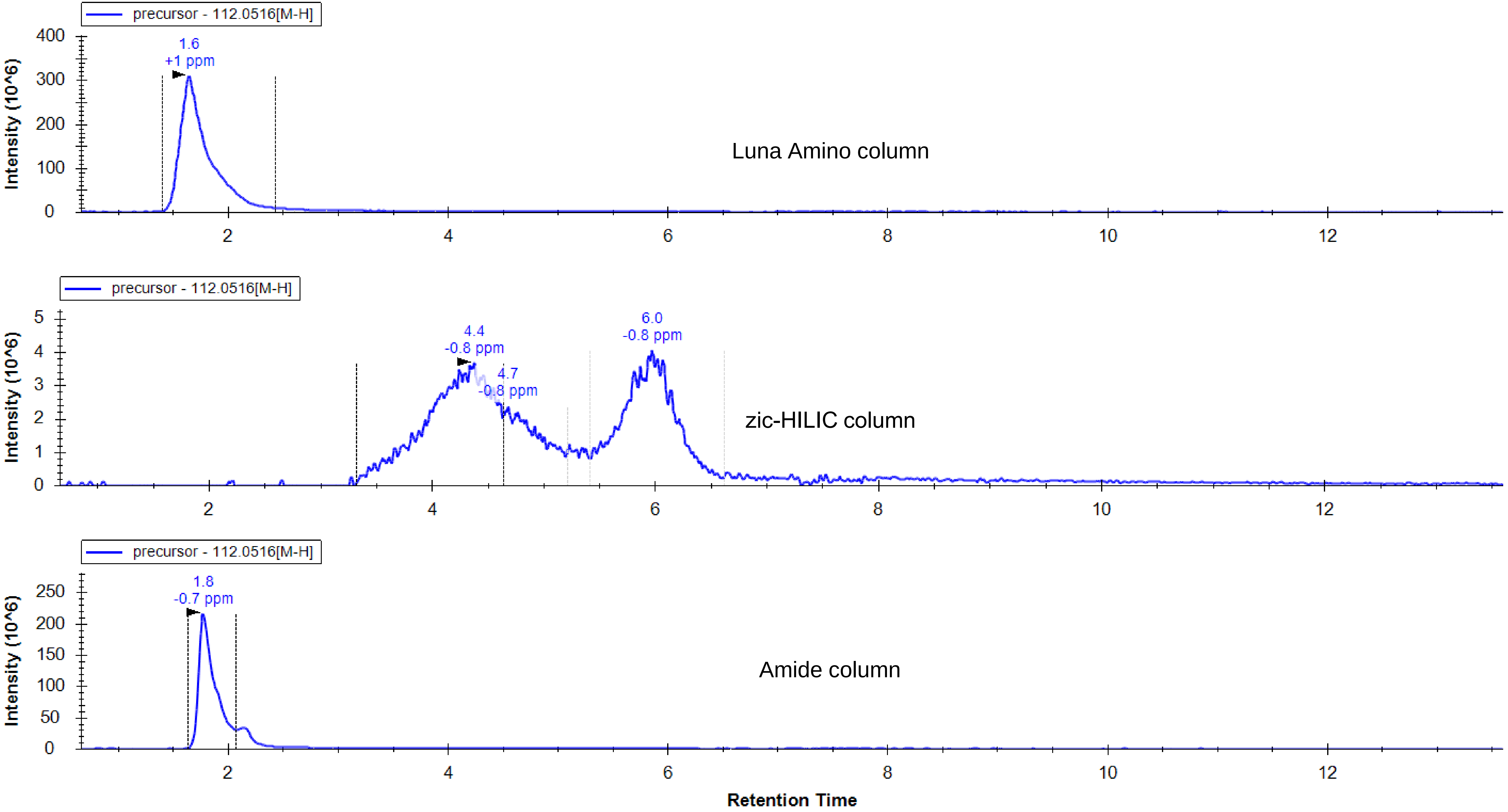

Supplementary Figure 10) Comparison of chromatographic performance of 3 columns for analysis of Tryptophan standard

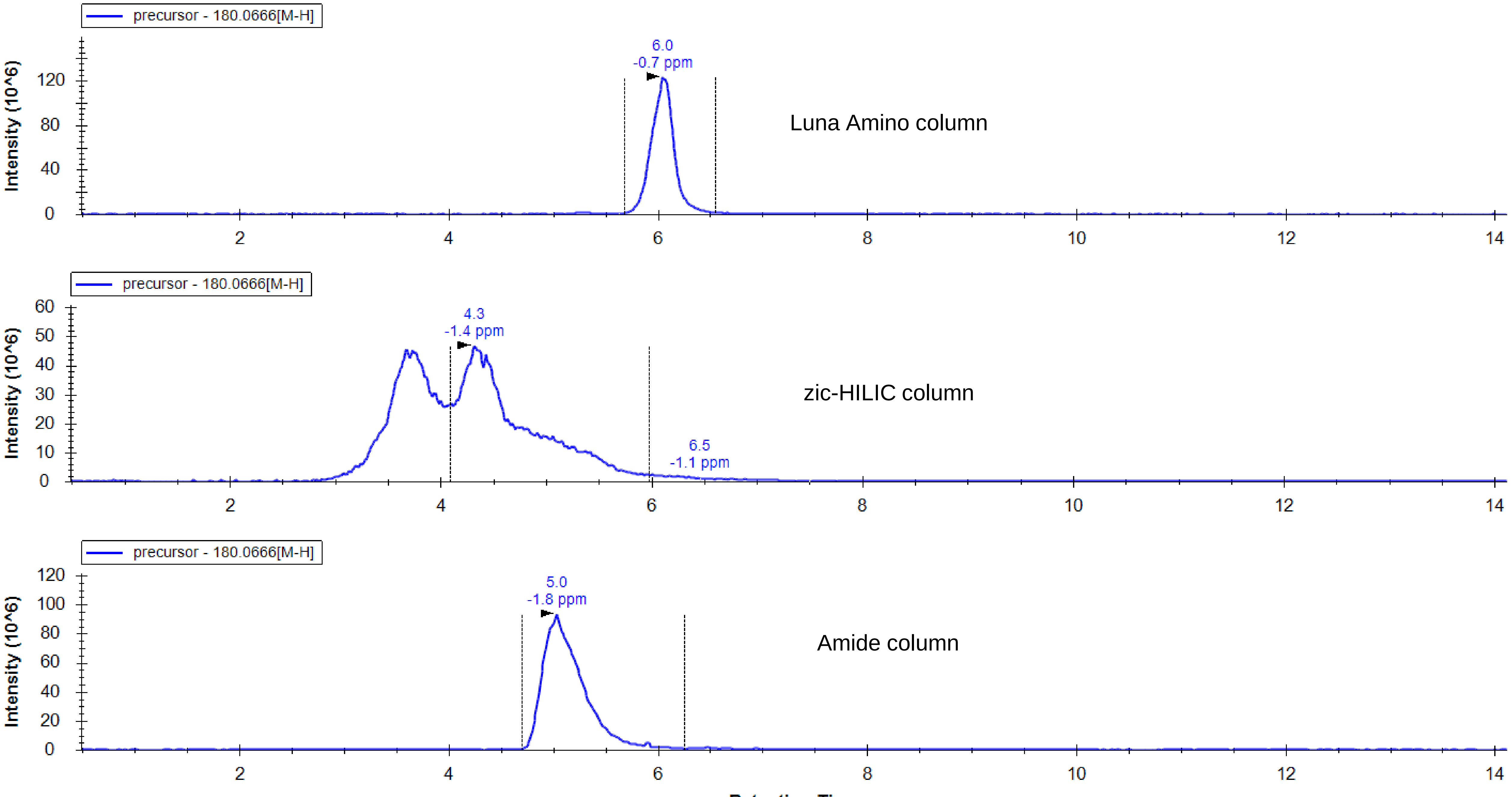

Supplementary Figure 11) Comparison of chromatographic performance of 3 columns for analysis of Carnosine standard

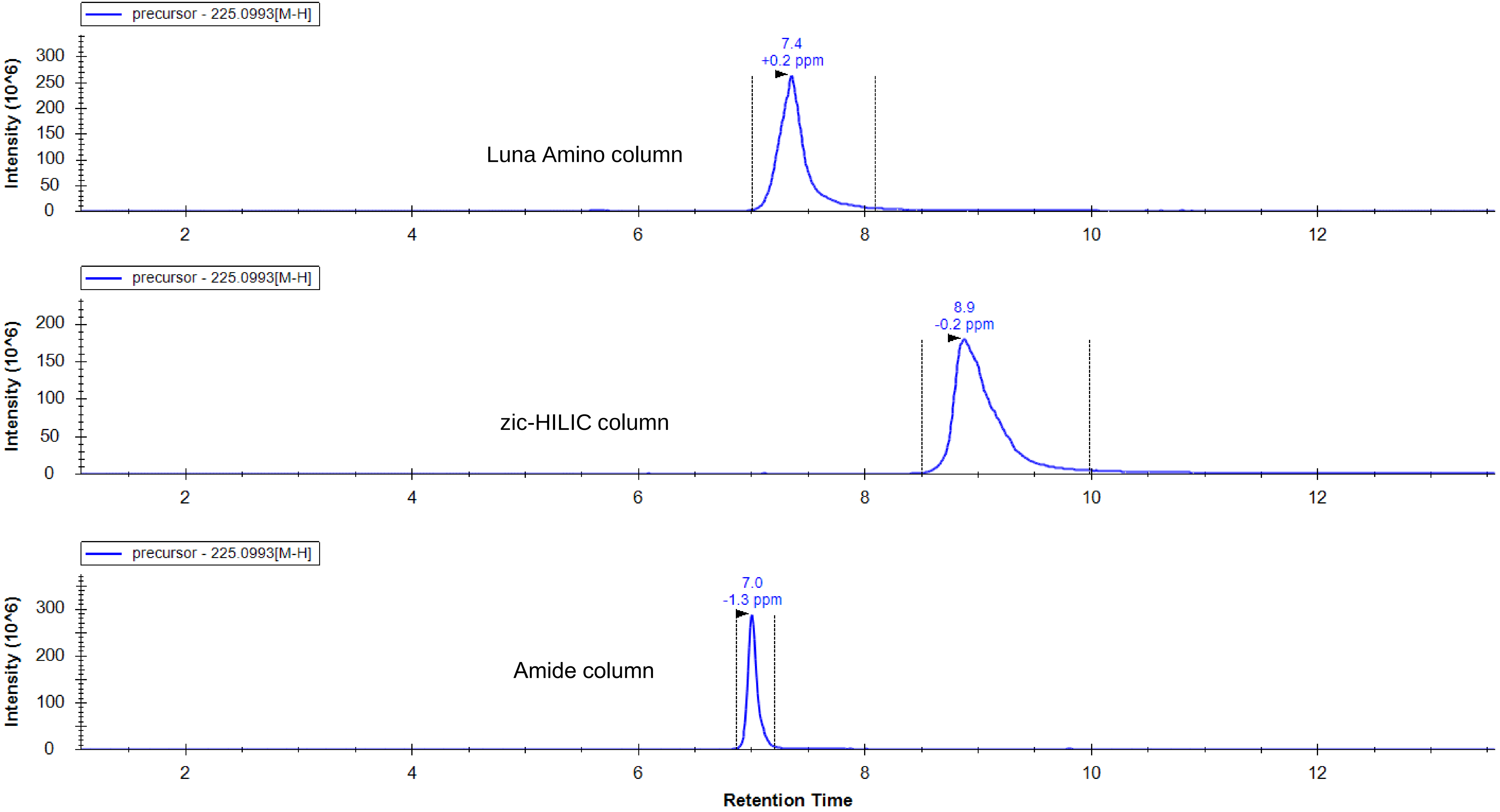

Supplementary Figure 12) Comparison of chromatographic performance of 3 columns for analysis of Glutamic acid standard

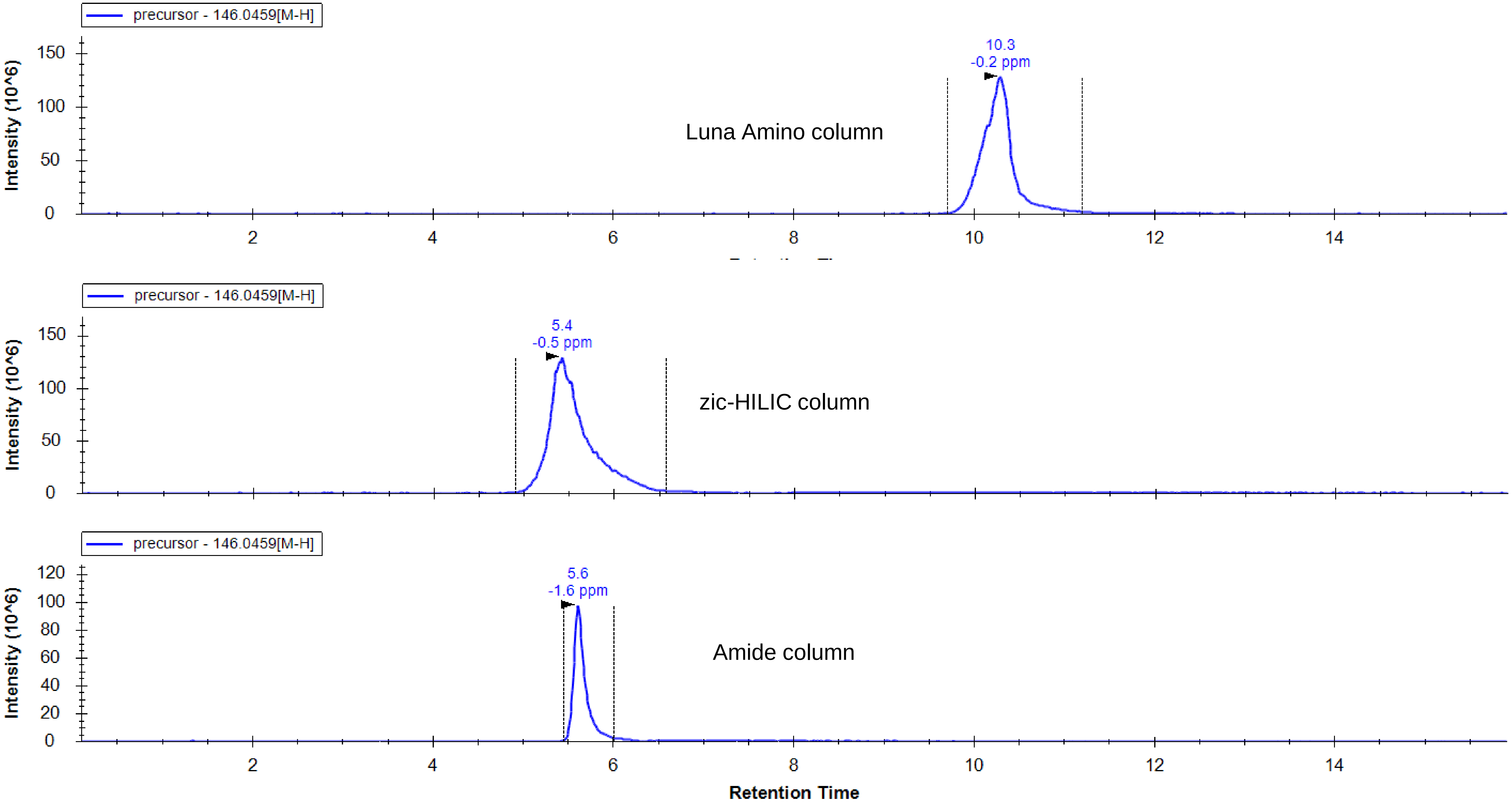
